# A Long Lost Key Opens an Ancient Lock: *Drosophila* Myb Causes a Synthetic Multivulval Phenotype in Nematodes

**DOI:** 10.1101/788265

**Authors:** Paul J. Vorster, Paul Goetsch, Tilini U. Wijeratne, Keelan Z. Guiley, Laura Andrejka, Sarvind Tripathi, Braden J. Larson, Seth M. Rubin, Susan Strome, Joseph S. Lipsick

## Abstract

The five-protein MuvB core complex (LIN9/Mip130, LIN37/Mip40, LIN52, LIN54/Mip120, and LIN53/p55CAF1/RBBP4) has been highly conserved during the evolution of animals. This nuclear complex interacts with proteins encoded by the *RB* tumor suppressor gene family and its associated E2F-DP transcription factors to form DREAM complexes that repress the expression of genes that regulate cell cycle progression and cell fate. The MuvB core complex also interacts with proteins encoded by the *Myb* oncogene family to form the Myb-MuvB complexes that activate many of the same target genes. We show that animal-type *Myb* genes and proteins are present in Bilateria, Cnidaria, and Placozoa, the latter including some of the simplest known animal species. However, bilaterian nematode worms appear to have lost their animal-type *Myb* genes hundreds of millions of years ago. Nevertheless, the amino acids in the LIN9 and LIN52 proteins that directly interact with the MuvB-binding domains of human B-Myb and *Drosophila* Myb are conserved in *C. elegans*. Here we show that, despite greater than 500 million years since their last common ancestor, the *Drosophila melanogaster* Myb protein can bind to the nematode LIN9 and LIN52 family proteins *in vitro* and can cause a synthetic multivulval (synMuv) phenotype *in vivo*. This phenotype is similar to that caused by loss-of-function mutations in *C. elegans* synMuvB class genes including those that encode homologs of the MuvB core, RB, E2F, and DP. Furthermore, amino acid substitutions in the MuvB-binding domain of *Drosophila* Myb that disrupt its functions *in vitro* and *in vivo* also disrupt its activity in *C. elegans*. We speculate that nematodes and other animals may contain another protein that can bind to LIN9 and LIN52 in order to activate transcription of genes repressed by DREAM complexes.

## INTRODUCTION

The *Myb* gene family was discovered due to the retroviral transduction of the *c-Myb* proto-oncogene that created the *v-Myb* oncogene of the avian myeloblastosis virus ^1^. Vertebrate animals including humans have three paralogous *Myb* genes (*A-Myb/MYBL1*, *B-Myb/MYBL2*, and *c-Myb/MYB*) ^2,3^. The fruit fly *Drosophila melanogaster* and many other invertebrate species contain a single essential animal-type *Myb* gene that is most closely related to vertebrate *B-Myb* ^2–7^. The vertebrate *A-Myb* and *c-Myb* genes appear to have arisen by two rounds of gene duplication and divergence from a *B-Myb*-like ancestral gene ^8^. Consistent with this model, vertebrate *B-Myb* but neither *A-Myb* nor *c-Myb* can complement the cell cycle defects observed in *Drosophila Myb* null mutant animals ^6,9^.

Animal Myb-type proteins all contain a broadly conserved pan-eukaryotic amino-terminal DNA-binding domain and animal-specific domains ^3^. The vertebrate A-Myb and c-Myb proteins also share a central transcriptional activation domain that is not well-conserved in vertebrate B-Myb or invertebrate Myb proteins. Surprisingly, the animal-specific carboxy-terminus of *Drosophila* Myb is both necessary and sufficient for rescue of the adult lethality of a *Myb* null mutant, for proper association with chromatin, for transcriptional activation of essential G2/M phase genes, and for mitotic cell cycle progression ^10,11^. Alanine substitutions of evolutionarily conserved motifs identified a short peptide sequence that is required for all these functions ^11^.

Biochemical purification of an activity that bound to DNA near a developmentally regulated origin of replication in a *Drosophila* chorion locus led to the discovery of a multiprotein complex that contains Myb and several Myb-interacting proteins (Mip) ^12^. Similar complexes called Myb-MuvB, which contain either B-Myb or less frequently A-Myb, were later identified in human cells ^13^. In addition to *Drosophila* Myb or vertebrate B-Myb, these complexes contain Mip130/LIN9, Mip120/LIN54, Mip40/LIN37, p55CAF1/RbAp48, and LIN52. These five additional proteins, known as the MuvB core, can also associate with *Drosophila* E2F2, DP, and RBF1 or RBF2 or their vertebrate homologs (E2F4 or E2F5, DP1 or DP2, p107 or p130) to form complexes now called DREAM ^13^. A large holocomplex containing Myb, E2F, DP, and RB family proteins together with the MuvB core and called dREAM was identified in *Drosophila* embryos, but has not been observed in human cell lines ^13–15^.

*Myb* loss-of-function mutants in *Drosophila* display mitotic cell cycle defects and aberrations in ploidy in somatic tissues ^6,7,16–19^. Both in *Drosophila* and in human cell lines, the Myb-MuvB complex was shown to activate the transcription of genes essential for G2/M progression in mitotically active cells, whereas the DREAM complex represses these genes ^20–26^. These complexes have also been implicated in human cancer initiation and progression. For example, a high level of *B-Myb/MYBL2* expression in breast cancer is a clinically useful predictor of tumor recurrence and decreased patient survival ^27–29^. Furthermore, extensive DNA sequencing has revealed that approximately one-half of human breast cancer specimens contain a genetic alteration in at least one of the genes encoding subunits of these two complexes (Figure S1).

Remarkably, all of the proteins in the Myb-MuvB and DREAM complexes, with the exception of Myb itself, are encoded by homologs of synMuvB group genes in the nematode *Caenorhabditis elegans* ^30^. In brief, dominant gain-of-function mutations in the EGF=>RAS=>RAF=>MEK=>MAPK=>ETS pathway caused a multivulval (Muv) phenotype in *C. elegans* ^31^. Recessive loss-of-function mutations in *lin-8* and *lin-9* together caused a synthetic multivulval phenotype (synMuv) ^32^. Additional genetic screens identified two groups of genes (synMuvA and synMuvB), in which any group A mutation could cooperate with any group B mutation to cause this synMuv phenotype ^33^. The proteins encoded by synMuvA and synMuvB genes redundantly repress ectopic expression of the secreted LIN-3/EGF protein that normally controls vulval development via the RAS pathway ^31,34,35^. The synMuvB genes also regulate transgene silencing, cell cycle progression, repression of germline-specific genes in somatic cells, RNA interference (RNAi), and X chromosome gene expression ^36–40^.

In *Drosophila*, the DREAM complex encoded by homologs of nematode synMuvB genes represses the expression of G2/M phase genes and also represses ectopic expression of the carbon dioxide receptor in olfactory neurons ^10,19,41^. The *Drosophila* Myb protein is required to relieve this DREAM-mediated repression for mitotic cell cycle progression and for carbon dioxide receptor expression in the appropriate neurons. *Drosophila* Myb also acts in opposition to the DREAM complex to regulate chorion gene amplification in ovarian follicle cells and programmed neuronal cell death ^12,42,43^. Interestingly, recent studies in *C. elegans* have shown that the MuvB complex can effectively repress gene expression in the absence of the LIN-35 RB-family protein that was previously thought to be required for repression by DREAM complexes ^44^. Although *C. elegans* and other nematode species contain two *Myb*-related genes that encode homologs of the CDC5/CEF1 splicing factor and the SNAPc small nuclear RNA transcription factor, they do not contain an animal-type *Myb* gene that might relieve repression by DREAM complexes ^3^.

The animal-specific carboxy-terminus of *Drosophila* Myb is both necessary and sufficient for binding to the MuvB core complex in cell lysates ^11^. In addition, two alanine substitution mutants that greatly diminished the biological activities of *Drosophila* Myb *in vivo* also inhibit its binding to the MuvB core complex. Studies with recombinant proteins identified conserved Myb-binding domains of human LIN9 and LIN52 that in concert are sufficient for binding to the homologous MuvB-binding domain of human B-Myb and *Drosophila* Myb ^45^. Structural determination by X-ray crystallography revealed a coiled-coil comprised of human LIN9 and LIN52 α-helices, which together form a binding site for the MuvB-binding domain of B-Myb. Furthermore, the amino acids in B-Myb homologous to those disrupted by the non-functional *Drosophila* Myb alanine substitution mutants make critical contacts with the LIN9 and LIN52 Myb-binding domains.

Surprisingly, the residues in human LIN9 and LIN52 that contact B-Myb are highly conserved in *C. elegans*, which itself lacks an animal-type Myb protein. Interestingly, in structure-based molecular modeling studies, the homologous peptides of nematode LIN9 and LIN52 could readily accommodate the MuvB-binding domains of animal Myb proteins. We therefore decided to experimentally test whether the MuvB-binding domain of *Drosophila* Myb can functionally interact with the putative Myb-binding domains of *C. elegans* LIN9 and LIN52, both *in vitro* and *in vivo*.

## RESULTS

### Evolutionary Conservation of Animal-Type Myb Proteins

Searches of public sequence repositories revealed that animal-type Myb proteins, characterized by an amino-terminal DNA-binding domain composed of three tandem Myb repeats, a central proline-rich “hinge”, and a carboxy-terminal MuvB-binding domain, are present in species of all phyla of the superphylum Deuterostomia, including Chordata (human, lancelet, sea squirt), Hemichordata (acorn worm), and Echinodermata (sea urchin) (Figures 1 and 2). Animal-type Myb proteins, defined as having these three domains, are also present in species of widely divergent invertebrate animal phyla including Arthropoda (fruit fly), Priapulida (penis worm), Mollusca (scallop), Brachiopoda (lamp shell), Cnidaria (coral), and Placozoa (*Trichoplax*). Surprisingly, none of the 25 completely sequenced and widely divergent species of Nematoda (round worm) contain an animal-type Myb protein, although they do contain more distant Myb-related proteins of the SNAPc and Cdc5/CEF1 families that are also present in a wide range of eukaryotes including fungi ^3^. Nematodes have not left a fossil record, but analyses of molecular evolution have led to estimates of between 600 and 1300 million years since their divergence from other animals ^46^.

**Figure 1.**
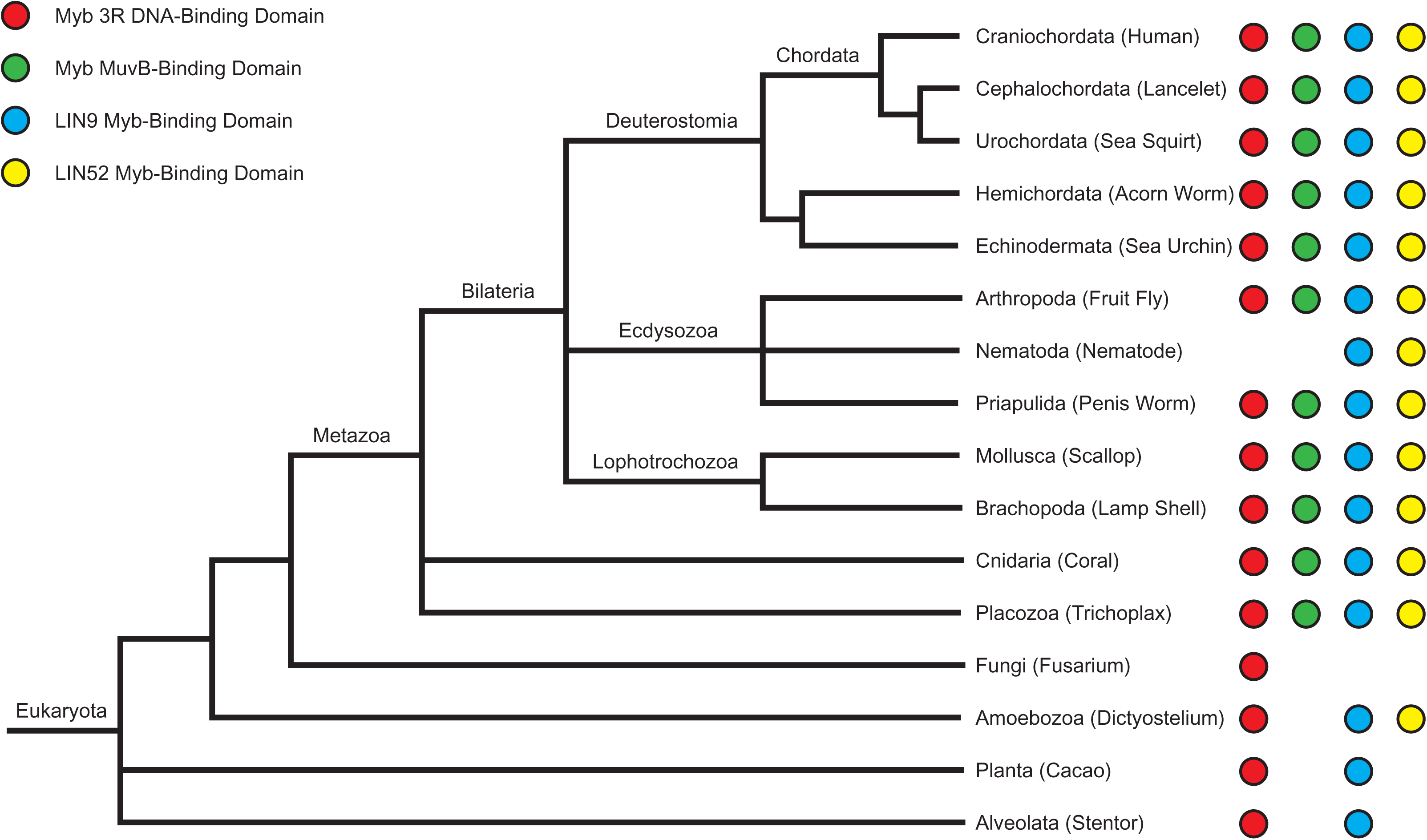
Evolutionary Conservation of Myb Three-Repeat (3R) DNA-Binding Domains, MuvB-Binding Domains of Myb Proteins, and Myb-Binding Domains of LIN9 and LIN52. A partial phylogenetic tree of the current view of eukaryotic evolution [http://tolweb.org/tree/] shows the presence or absence of the indicated protein domains in representative species from diverse clades: Human (*Homo sapiens*), Lancelet (*Branchiostoma belcheri*), Sea Squirt (*Ciona intestinalis*), Acorn Worm (*Saccoglossus kowalevskii*), Sea Urchin (*Strongylocentrotus purpuratus*), Fruit Fly (*Drosophila melanogaster*), Nematode (*Caenorhabditis elegans*), Penis Worm (*Priapulus caudatus*), Scallop (*Mizuhopecten yessoensis*), Lamp Shell (*Lingula anatina*), Coral (*Stylophora pistillata*), Trichoplax (*Trichoplax adhaerens*), Fusarium (*Fusarium sp. AF-4*), Dictyostelium (*Dictyostelium discoideum*), Cacao (*Theobroma cacao*), Stentor (*Stentor coeruleus*). Displayed branch lengths are unscaled.

**Figure 2.**
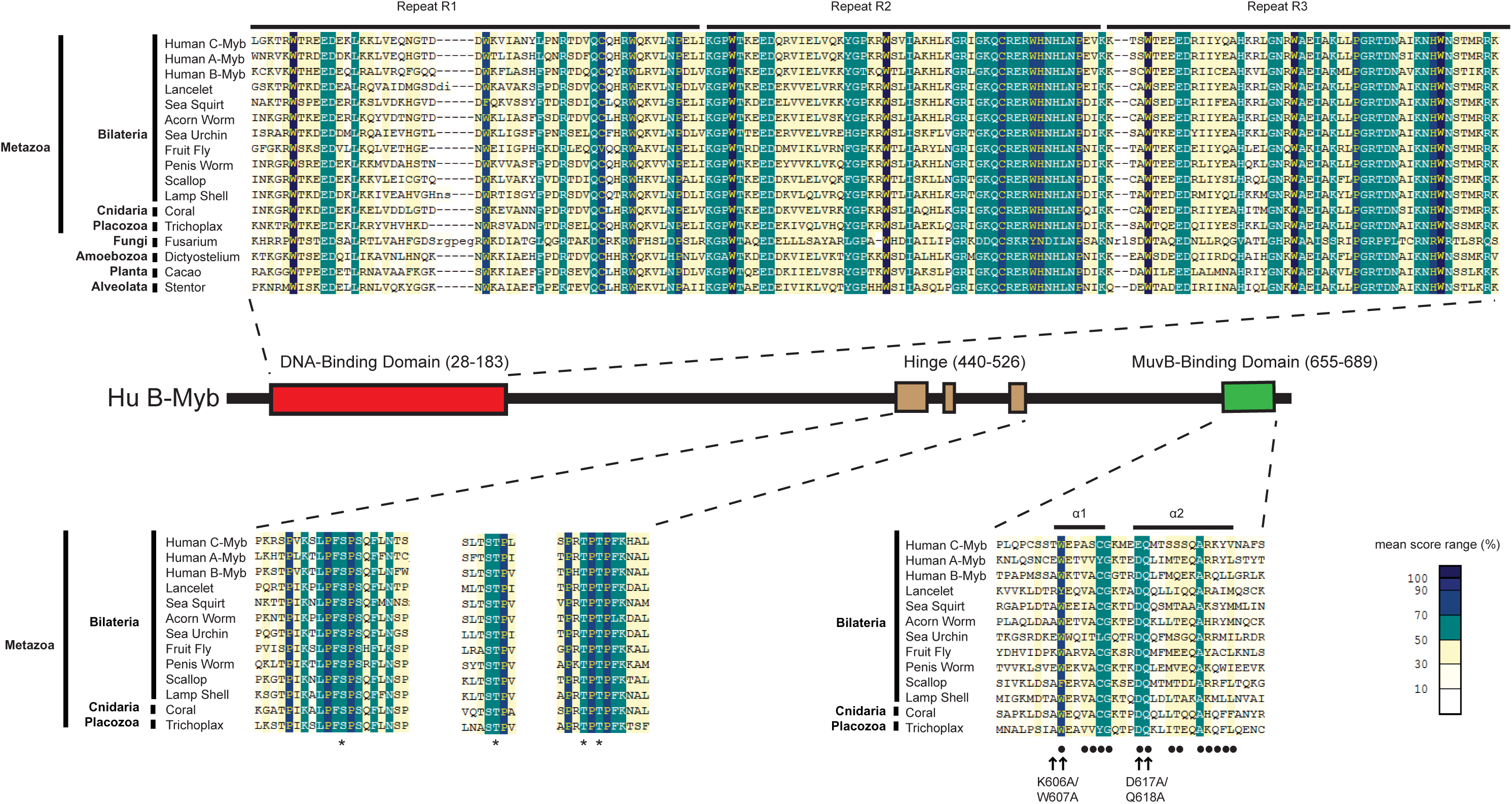
Sequence Alignments of Conserved Animal-Type Myb Protein Domains. A schematic diagram of the human B-Myb protein shows the relative positions and amino acid sequence numbers of the conserved domains that define animal-type Myb proteins. Local multiple protein sequence alignments were constructed using MACAW with the BLOSUM62 scoring matrix ^63^. The alignment shading indicates the mean score at each position as shown in the color key. Horizontal bars above the DNA-binding domain alignment indicate three tandem Myb repeats (R1, R2, R3). Asterisks below the hinge domain alignments indicate known Cyclin A-CDK2 phosphorylation sites in human B-Myb ^62^. The central hinge domain alignment contains a binding site for the Plk1 polo-family protein kinase. Horizontal bars above the MuvB-binding domain alignment indicate α-helices in the human B-Myb crystal structure with human LIN9 and LIN52 ^45^. Black dots below the MuvB-binding domain alignment indicate amino acids of human B-Myb that contact human LIN9 or LIN52 in the crystal structure. Arrows below the MuvB-binding domain indicate amino acids substituted by alanine in two *Drosophila* Myb mutants used in experiments in this study ^11^.

The phylogenetic relationship of nematodes to chordates and arthropods within the Bilateria has been controversial ^47,48^. Nevertheless, the presence of animal-type *Myb* genes in species of the non-Bilaterian “out group” phyla Cnidaria (coral) and Placozoa (*Trichoplax adherens*), argues strongly that animal-type *Myb* genes were present in the last common ancestor of all these widely divergent animal species, including nematodes. Therefore, a common ancestor of all modern nematodes appears to have lost its animal-type *Myb* gene. A similar loss may have occurred during the evolution of some other phyla of the Bilateria, but in those cases there are not as many divergent species with completely sequenced genomes as in the Nematoda.

The presence of highly conserved three-repeat Myb DNA-binding domains in species of the kingdoms of Fungi (*Fusarium*) and Plants (*Cacao*) and in the “orphan” clades of Amoebozoa (*Dictyostelium*) and Avleolata (*Stentor*) suggests that a common ancestor of most if not all modern eukaryotes contained this domain (Figure 1 and 2). However, the MuvB-binding domain and adjacent proline-rich hinge that are also present in animal-type Myb proteins have not been found in any of the known Myb-related proteins of widely divergent non-animal species. This result suggests that the MuvB-binding domain and adjacent proline-rich hinge emerged subsequent to the divergence of Metazoa from other eukaryotic kingdoms and “orphan” clades.

### Evolutionary Conservation of the Myb-Binding Domains of LIN9 and LIN52

Searches of public sequence repositories revealed that the Myb-binding domain of LIN9 proteins is conserved in a wide range of species within the Metazoa, including those in the Nematoda that lack an animal-type Myb protein (Figures 1 and 3). Sequences homologous to the Myb-binding domain of LIN9 were also identified in species within the Planta, Amoebozoa, and Alveolata that have proteins containing an animal-type Myb DNA-binding domain but lacking a MuvB-binding domain. The deep evolutionary conservation of the Myb-binding domain of LIN9 in the absence of the MuvB-binding domain of Myb suggests that the former domain is likely to have another function. The lack of any proteins homologous to either the DIRP (<u>d</u>omain in <u>R</u>b-related <u>p</u>athway; Pfam 06584) ^49^ or Myb-binding domains of LIN9 in Fungi suggests that a common ancestor of modern fungal species lost its *LIN9* gene after its divergence from the other kingdoms of eukaryotes. In addition, the presence of LIN9-related proteins with a DIRP domain but not a Myb-binding domain in some plant species including the intensively studied thale cress (*Arabidopsis thaliana*) suggests that these domains can function independently of one another.

**Figure 3.**
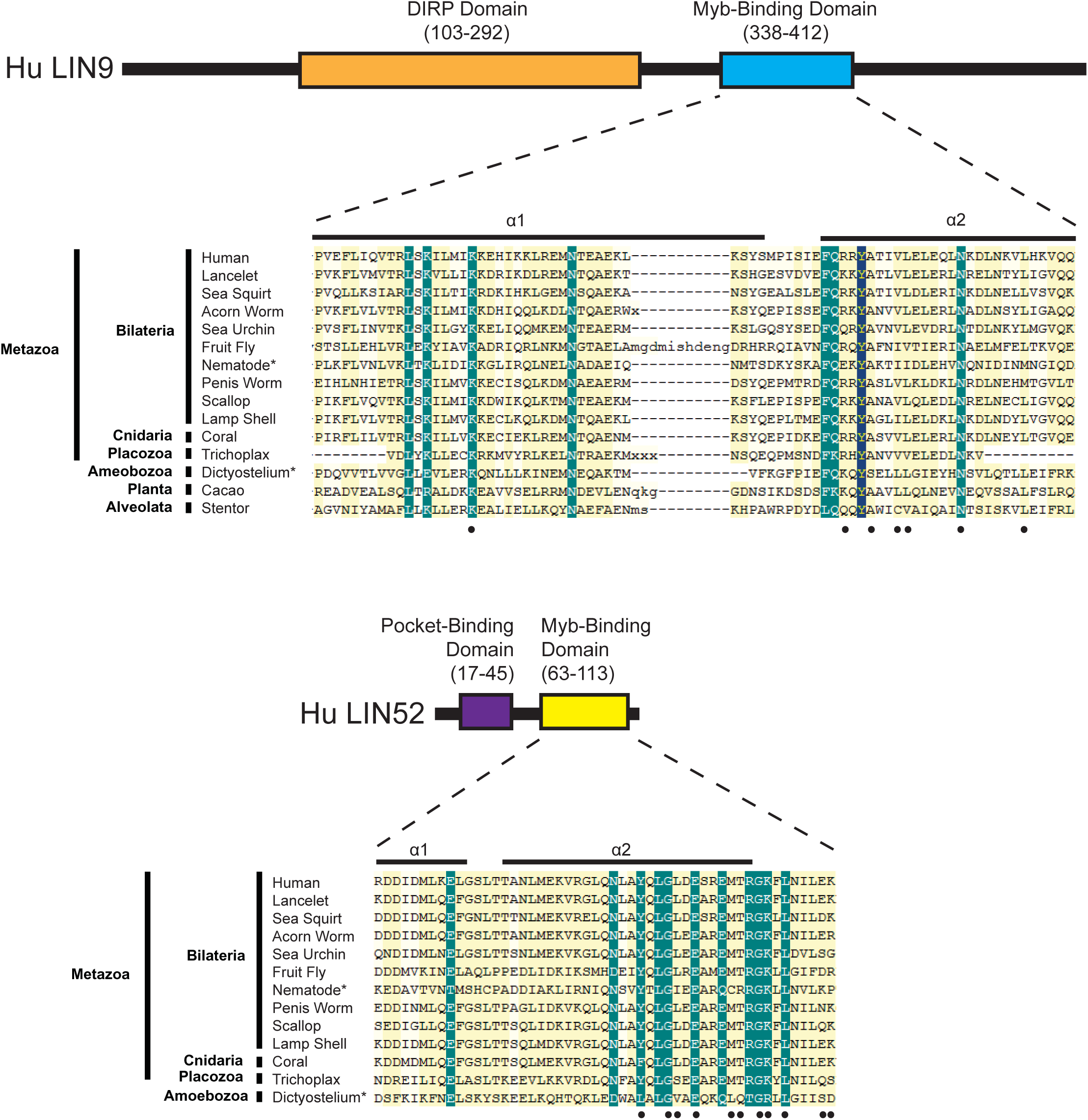
Sequence Alignments of the Myb-Binding Domains of LIN9 and LIN52. Schematic diagrams of the human LIN9 and LIN52 proteins show the relative positions and amino acid sequence numbers of the conserved domains. Horizontal bars above the Myb-binding domain alignments indicate α-helices in the crystal structure of human LIN9 and LIN52 bound to human B-Myb ^45^. Black dots below the MuvB-binding domain alignments indicate amino acids that contact human B-Myb in the crystal structure. Alignments are not shown for the DIRP domain of LIN9 (pfam 06584) ^49^, the function of which remains unknown, or for the pocket-binding domain of LIN52 that binds to the human RB-related p107 and p130 proteins but not to human RB itself ^66^.

Proteins of the LIN52 family are not as evolutionarily widespread as those of the LIN9 family and were only identified in species within the Metazoa and Amoebozoa (Figures 1 and 3). These LIN52 homologs contain both a pocket-binding domain and a Myb-binding domain. The former is consistent with the presence of RB “pocket” family proteins in these species. The absence of Myb family MuvB-binding domains in Nematoda and Amoebozoa, despite the presence of both LIN9 and LIN52 Myb-binding domains, again suggests that these domains have an additional evolutionarily conserved function.

### The MuvB-Binding Domain of Drosophila Myb Can Bind to a Nematode LIN9-LIN52

Although no sequenced species of Nematoda contains an animal-type Myb protein, they nevertheless do contain conserved Myb-binding domains in their LIN9 and LIN52 family proteins (Figures 1 and 3). Furthermore, the amino acids in the human LIN9 and LIN52 proteins that make direct contacts with human B-Myb in a structure determined by X-ray crystallography are well-conserved in the LIN9 and LIN52 proteins of the intensively studied nematode *Caenorhabditis elegans* ^45^ (Figure 3). Molecular modeling based on this structure predicts that the *C. elegans* LIN9 and LIN52 proteins are likely to be capable of binding to the MuvB-binding domain of *Drosophila* Myb (Figure 4). Furthermore, conserved amino acids in *Drosophila* Myb (K606, W607, D617, Q618) that were previously shown to mediate biochemical interactions with the *Drosophila* MuvB core complex *in vitro* and to be required for the function of *Drosophila* Myb *in vivo* are predicted to contact *C. elegans* LIN9 and LIN52 ^11^. The homologous amino acids in human B-Myb were shown to contact human LIN9 and LIN52 in the crystallographic structure ^45^.

**Figure 4.**
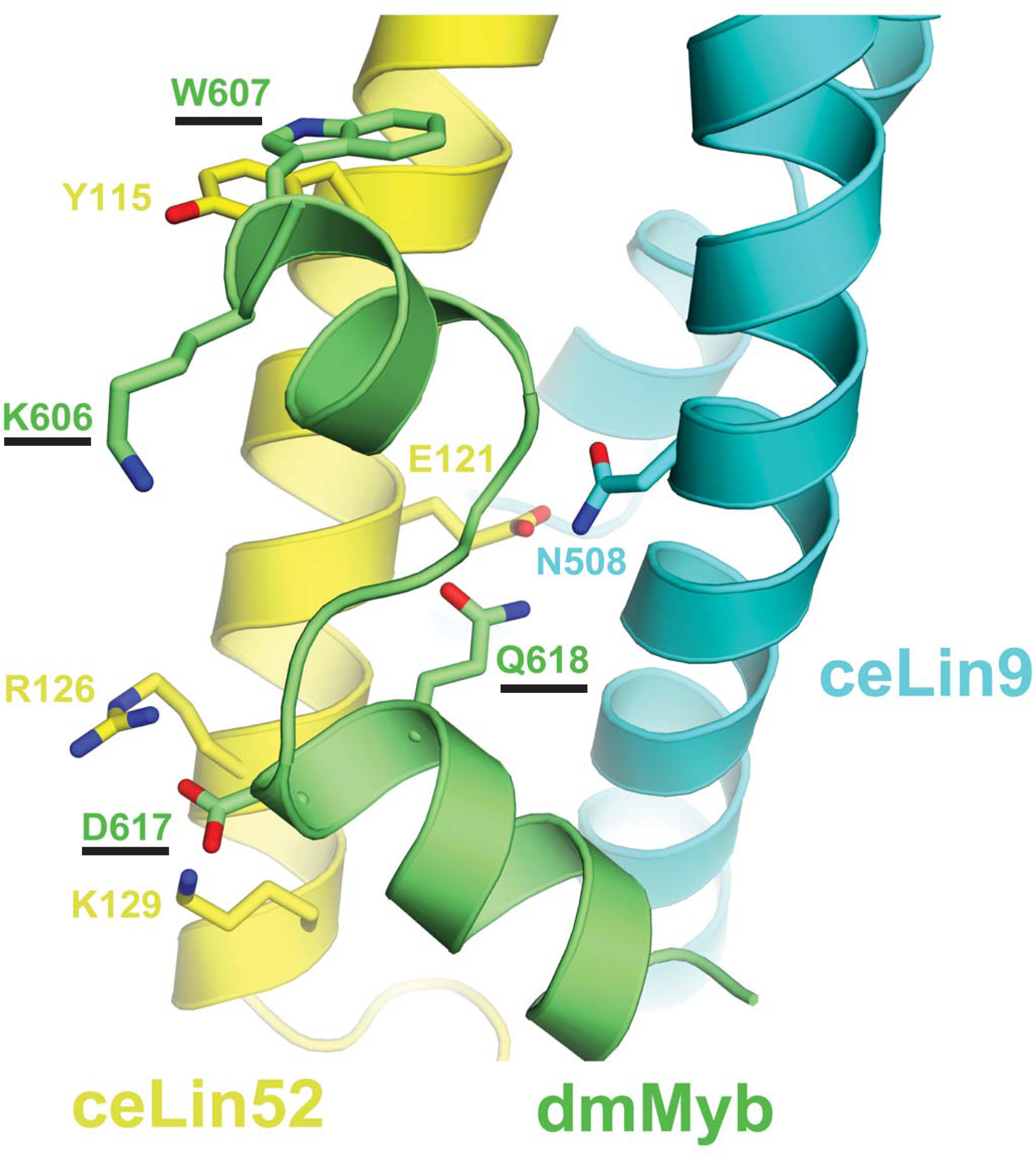
Structural Modeling of the *Drosophila* Myb MuvB-Binding Domain Bound to the Myb-Binding Domains of Nematode LIN9 and LIN52. Protein sequences of *Drosophila* Myb (dmMyb; green), *C. elegans* LIN9 (ceLIN9; cyan), and *C. elegans* LIN52 (ceLIN52; yellow) were modeled into the human crystal structure (PDB ID: 6C48) using MODELLER ^67^. The four amino acids substituted by alanine in the two *Drosophila* Myb mutants used in experiments in this study are underlined.

To test whether the conservation of Myb-binding domain sequence in *C. elegans* LIN9 and LIN52 results in conservation of protein function, the relevant recombinant protein domains were produced in *E. coli* and purified using affinity, ion-exchange, and size-exclusion chromatography (Figure S2). Reconstituted heterodimeric LIN9-LIN52 Myb-binding domain complexes were tested for their ability to bind the MuvB-binding domain of *Drosophila* Myb using isothermal titration calorimetry (Figure 5). The *C. elegans* and the *Drosophila* LIN9-LIN52 heterodimers bound to *Drosophila* Myb with very similar affinities (K_D_ = 7 or 3 μM, respectively). The K606A/W607A double substitution mutant of *Drosophila* Myb caused an approximately 10-fold reduction of binding to either *C. elegans* or *Drosophila* LIN9-LIN52 (K_D_ = 70 or 41 μM, respectively). Furthermore, the D617A/Q618A double substitution mutant of *Drosophila* Myb abolished any detectable binding to either *C. elegans* or *Drosophila* LIN9-LIN52. These experiments show that despite over 500 million years since nematodes appear to have lost their animal-type Myb genes, their LIN9 and LIN52 proteins are still capable of binding to the MuvB-binding domain of *Drosophila* Myb in a similar fashion to *Drosophila* LIN9 and LIN52.

**Figure 5.**
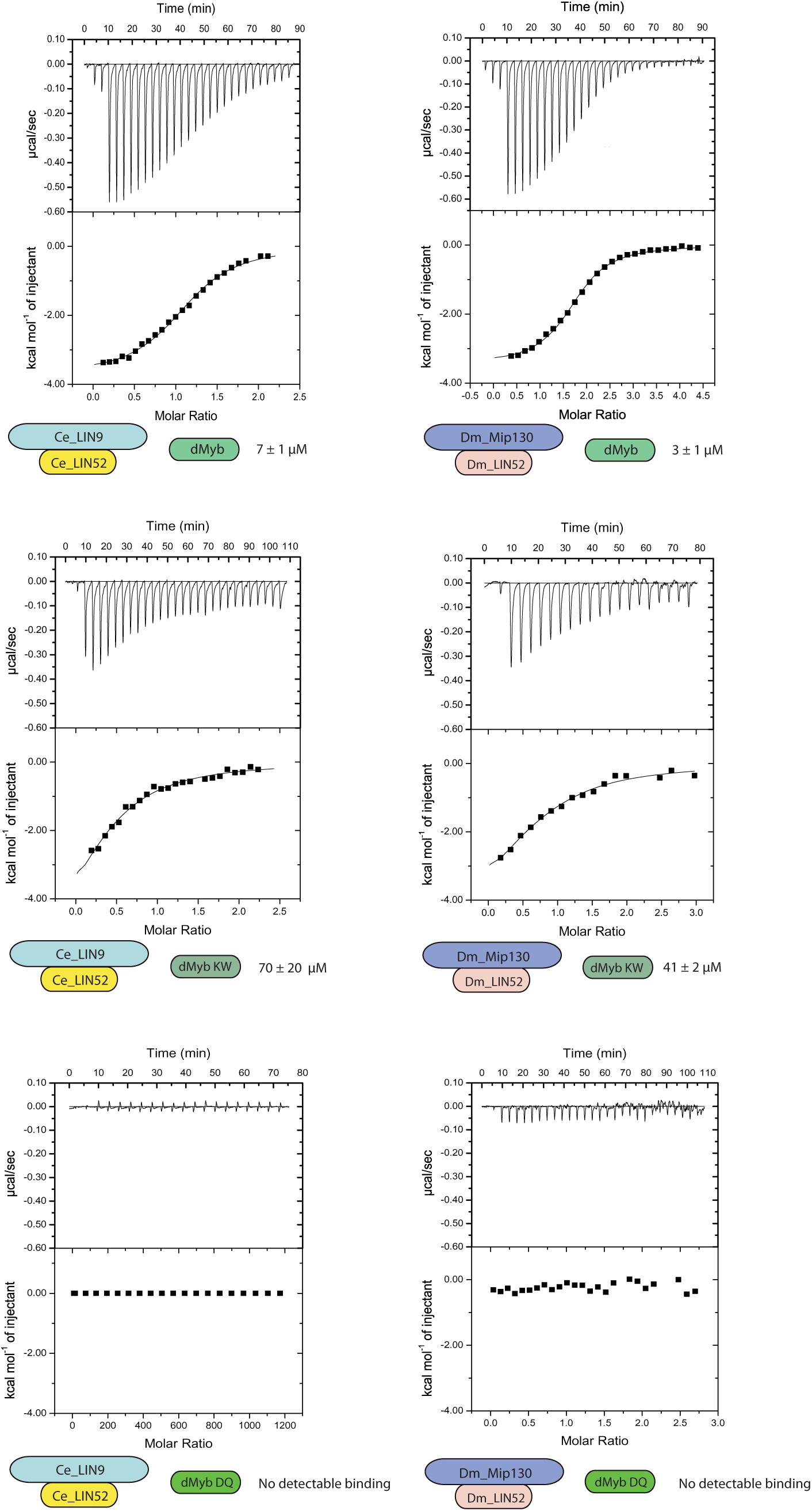
The Myb-Binding Domains of *C. elegans* LIN9 and LIN52 Bind the *Drosophila* MuvB-Binding Domain of *Drosophila* Myb *In Vitro*. Recombinant *C. elegans* LIN9-LIN52 heterodimeric Myb-binding domains produced in *E. coli* were purified and then assayed for binding to recombinant *Drosophila* Myb MuvB-binding domain using isothermal titration calorimetry. Each panel displays a representative experiment using the proteins diagrammed below the panel. The raw data are presented above and the fitted binding curve is presented below. The mean calculated K_D_ values and standard deviations from three replicate experiments are shown below each panel.

### Expression of Drosophila Myb in C. elegans Causes a Synthetic Multivulval Phenotype

The *lin-9* gene was discovered in *C. elegans* because a loss-of-function mutation cooperated with a second loss-of-function mutation in *lin-8* to cause a synthetic multivulval (synMuv) phenotype ^32^. Additional genetic screens identified a group of genes (class A synMuv) for which a loss-of-function mutant could cooperate with a *lin-9* mutant to cause a synMuv phenotype. A second group of genes (class B synMuv) including both *lin-9* and *lin-52* were identified for which a loss-of-function mutant could cooperate with a class A synMuv loss-of-function mutant to cause a synMuv phenotype ^33^. Since the *Drosophila* Myb protein can activate genes *in vivo* that are repressed by homologs of proteins encoded by synMuvB genes including Mip130/LIN9, E2F2, RBF1, and RBF2, we hypothesized that expression of *Drosophila* Myb in *C. elegans* might cause a synMuv phenotype similar to that seen in loss-of-function mutants of the endogenous synMuvB genes ^10,19,41^.

To test this hypothesis, GFP::Myb fusion proteins or a GFP-only (GFP) control protein were expressed in *C. elegans* under control of a heat-shock promoter using stably integrated single-copy transgenes. This was accomplished using the CRISPR-Cas9 system to promote transgene integration at a specific site on chromosome II ^50,51^. Four different GFP::Myb fusion proteins were expressed in this manner: full-length wild type (Myb), a mutant of Myb lacking its N-terminal DNA-binding domain but containing its C-terminal MuvB-binding domain (C-term), the K606A/W607A (KW) double substitution mutant, and the D617A/Q618A (DQ) double substitution mutant. Homozygous transgenic strains were genotyped by PCR of genomic DNA (Figure S3). Following heat shock of transgenic worms, nuclear GFP fluorescence following induction by heat shock. Following heat shock of transgenic worms, GFP expression was detected in nuclei in the majority of animals examined. Bright nuclear fluorescence in the polyploid gut cells was readily detected using a dissecting microscope with a 20X objective (Figure S4). Detailed examination via live spinning disc confocal microscopy with at 40X objective revealed nuclear GFP expression in the smaller vulval precursor cells (VPCs) and adjacent hypodermal cells (Figure 4S). Transgene expression varied from animal to animal, and from cell to cell within individual animals.

The transgenes were each crossed into a strong synMuvA (*lin-15A*) loss-of-function mutant background and doubly homozygous strains were isolated. These new strains were examined for the adult synMuv phenotype following induction of transgenic GFP or GFP::Myb protein expression by a single 15-minute heat shock during the L2/L3 stages of larval development. Consistent with a previous report of temperature-dependence, the *lin-15A* allele we used caused a Muv phenotype in approximately 5% of worms following heat shock with or without the *GFP* control transgene ^35^. In contrast, induction of the full-length wild type *GFP::Myb* transgene in the same *lin-15A* mutant background elevated the incidence of a synMuv phenotype to approximately 20% of worms (Figure 6).

**Figure 6.**
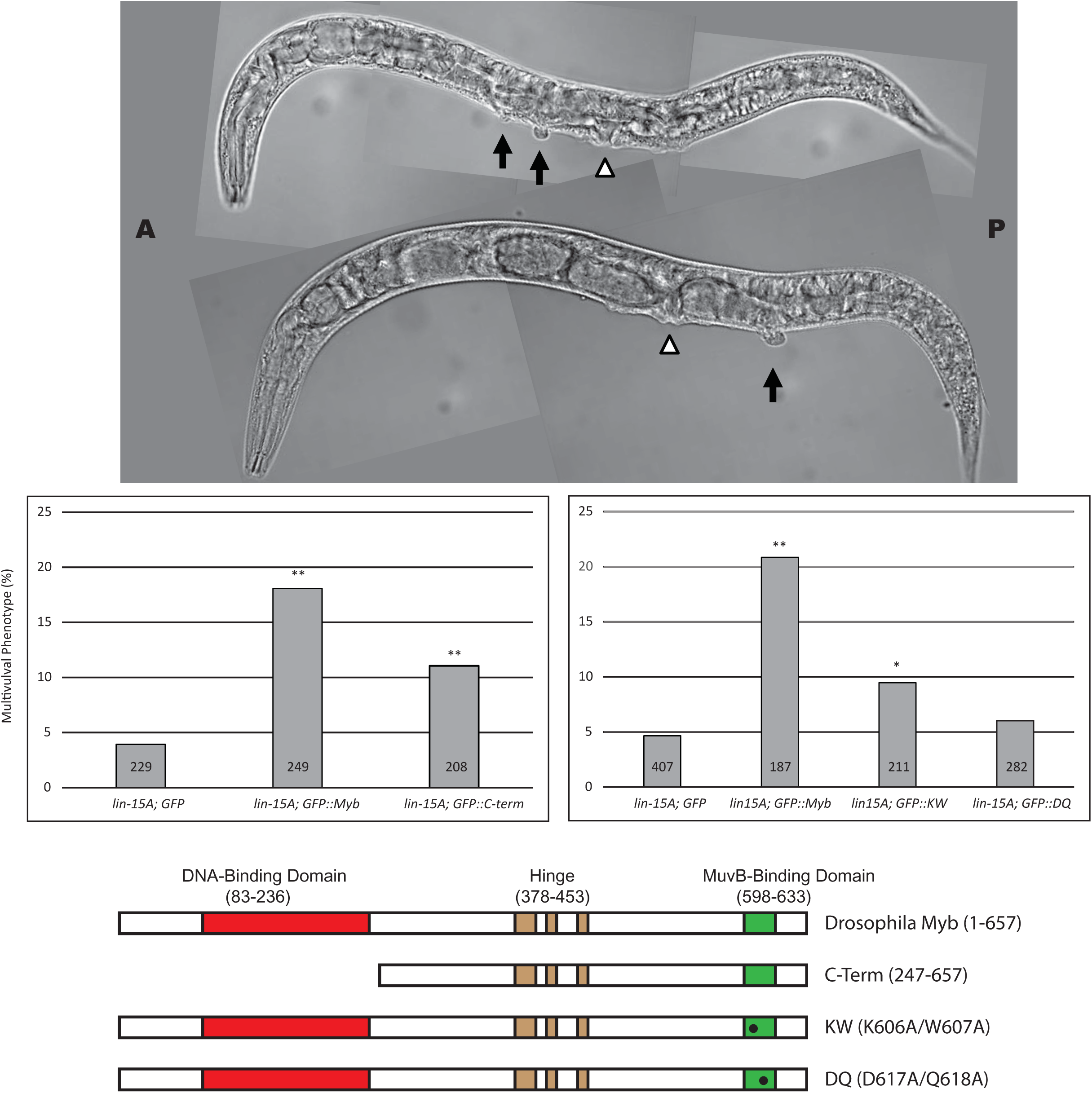
*Drosophila* Myb Causes a Synthetic Multivulval Phenotype in *C. elegans*. Top panel: The indicated strains were heat-shocked as L2/L3 larvae, then scored as adults for the presence of a multivulval phenotype in a *lin-15A* mutant background. DIC images of two representative multivulval worms of the *lin-15A; GFP::Myb* genotype are shown. The open arrowheads indicate the normal vulval opening. The black arrows indicate ectopic vulvae. Middle panel: The histograms show the incidence of multivulval worms in two different experiments using strains of the indicated genotypes. Statistical significance relative to the *lin-15A: GFP* control strain was determined using a two-tailed Z test. One asterisk indicates a significance of 0.05 or less; two asterisks indicate a significance of 0.01 or less. Numbers within the bars indicated total number of worms scored for the indicated genotype. Bottom panel: Schematic diagrams of the Drosophila Myb wild-type and mutant proteins expressed in transgenic worms. All of the Myb proteins contained a GFP tag fused at their amino termini (not shown).

A mutant of *Drosophila* Myb protein lacking its DNA-binding domain (C-term) was previously shown to be capable of physically interacting with the *Drosophila* MuvB core complex ^11^. Furthermore, in *Myb* null mutant flies, the C-term Myb mutant protein localizes to the cell nucleus, localizes to chromatin, activates the expression of G2/M phase genes, rescues G2/M phase cell cycle progression, and rescues adult viability at low temperatures ^10,11^. The C-term *Drosophila* Myb mutant protein also caused an elevated incidence of the synMuv phenotype when in combination with a *lin-15A* mutation, albeit less efficiently (approximately 10% incidence) than the full-length Myb protein (approximately 20% incidence) (Figure 6). Neither *Drosophila* Myb protein caused a multivulval phenotype in worms that were wild-type for *lin-15A*. These results show that the *Drosophila* Myb protein in combination with loss of *lin-15A* causes a synMuv phenotype in *C. elegans*, which itself has no animal-type Myb gene or protein of its own Furthermore, as was previously observed in *Drosophila*, the highly conserved Myb DNA-binding domain is not required for this synMuv phenotype in *C. elegans*.

The KW and DQ double alanine substitution mutants of *Drosophila* Myb were previously shown to abolish detectable immunoprecipitation of the *Drosophila* MuvB core complex from cell extracts *in vitro* and to be greatly diminished in rescuing expression of G2/M phase genes, cell cycle progression, and adult viability *in vivo* in *Myb* null mutant flies ^11^. Similar mutants of the human B-Myb protein were subsequently shown to inhibit its physical interaction with the human LIN9-LIN52 Myb-binding domain complex ^45^. The KW mutant of *Drosophila* Myb resembled the C-terminal Myb transgene in causing an approximately 10% incidence of the synMuv phenotype when in combination with a *lin-15A* mutation *Drosophila* (Figure 6). The DQ mutant of *Drosophila* Myb did not cause an elevation in the incidence of a synMuv phenotype in a *lin-15A* mutant background relative to the controls (all approximately 5%).

The differing abilities of the KW and DQ mutants of *Drosophila* Myb to cause a synMuv phenotype in *C. elegans* correlate with their abilities to bind either weakly or not at all to the *C. elegans* LIN9-LIN52 Myb-binding domain complex *in vitro* (Figure 5). The relative activities of these mutant proteins in nematodes also correlate with the ability to rescue the adult viability of *Drosophila Myb* null mutants at low temperature under control of the *Myb* promoter: the KW mutant weakly rescues, while the DQ mutant does not rescue ^11^. Taken together these genetic and biochemical results with mutant Myb proteins suggest that the *C. elegans* LIN9 and LIN52 Myb-binding domains interact with the MuvB-binding domain of *Drosophila* Myb in a fashion very similar to that predicted by evolutionary conservation of protein sequences (Figures 2 and 3) and by structural homology modeling (Figure 4).

## DISCUSSION

The MuvB-binding domain of animal-type Myb proteins and the Myb-binding domains of the MuvB subunits LIN9 and LIN52 are conserved in diverse clades of modern Metazoa. Nematodes appear to have lost their animal-type Myb genes and proteins after their divergence from other clades of modern Metazoa (Figures 1 and 2). Nevertheless, the Myb-binding domains of nematode LIN9 and LIN52 have been highly conserved over more than 500 million years in the absence of Myb (Figures 1 and 3). Remarkably, the LIN9-LIN52 Myb-binding domain of *C. elegans* can still bind the MuvB-binding domain of *Drosophila* Myb *in vitro* with a similar affinity and discrimination between mutants as the homologous LIN9-LIN52 domain of *Drosophila* (Figures 4 and 5). Furthermore, the reintroduction of *Drosophila* Myb into *C. elegans* in a synMuvA mutant background caused a synthetic multivulval phenotype similar to that caused by synMuvB mutants (Figures 6). These results imply that Myb can act in opposition to transcriptional repression by DREAM-related complexes in *C. elegans*, just as it does in *Drosophila* and in human cell lines (Figure 7) ^10,22,26^.

**Figure 7.**
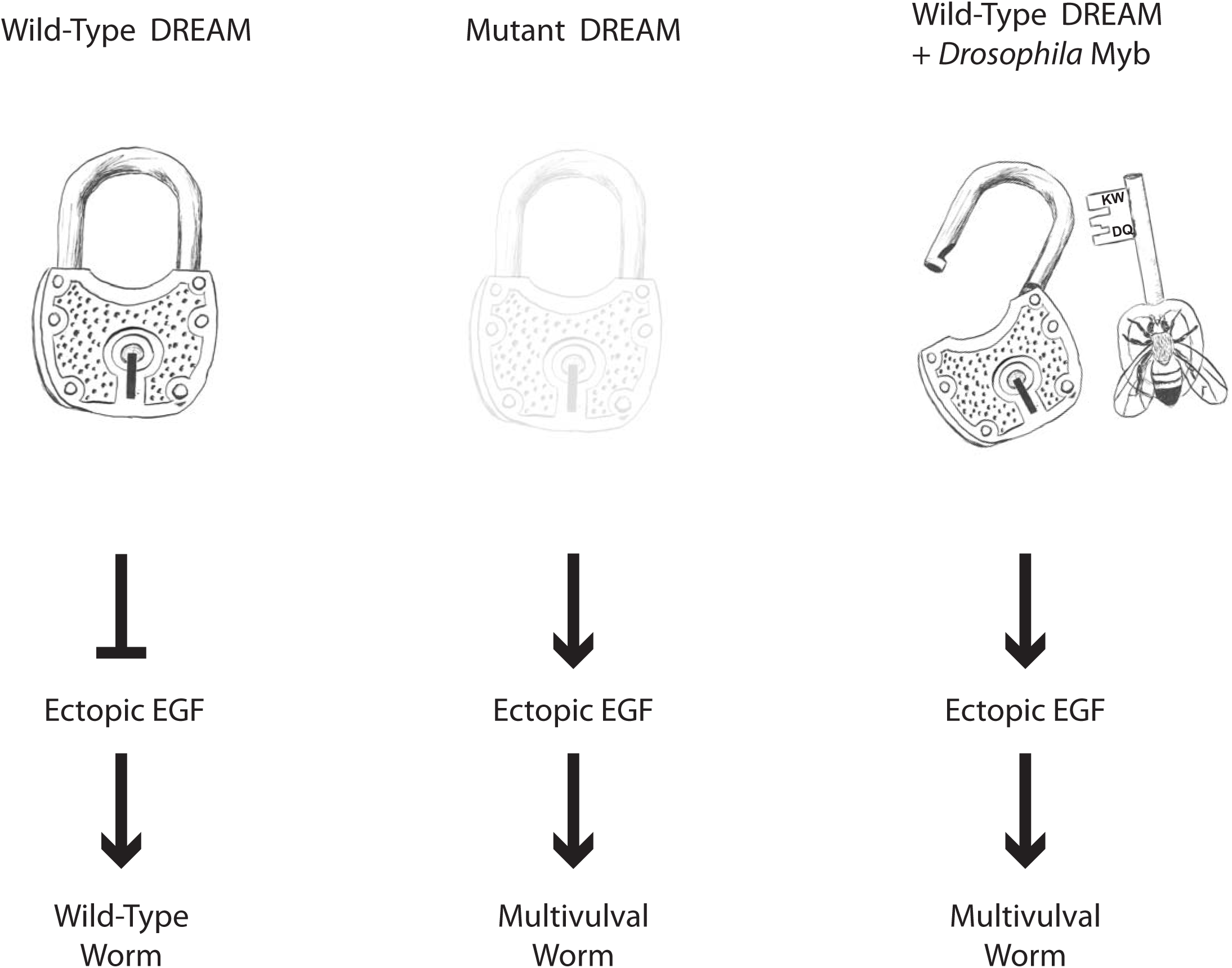
Model for the Mechanism of Action of *Drosophila* Myb in *C. elegans*. The wild-type DREAM complex, which includes LIN9 and LIN52 and other synMuvB proteins, redundantly represses the ectopic expression of LIN3/EGF, resulting in a wild-type worm even in a *lin-15A* synMuvA mutant background. Loss-of-function mutants of synMuvB genes fail to repress ectopic expression of LIN3/EGF, resulting in a synthetic multivulal worm in a *lin-15A* mutant background. Ectopic expression of *Drosophila* Myb overrides repression by the wild-type DREAM complex, causing a synthetic multivulval worm in a *lin-15A* mutant background, presumably due to ectopic expression of LIN-3/EGF. It remains unknown whether nematodes and other animals have a second “key” that can also open the highly conserved DREAM complex “lock”.

It will be interesting to determine whether ectopic expression of *Drosophila* Myb can mimic other phenotypes caused by synMuvB loss-of-function mutants in *C. elegans*. In this regard, we did not observe the misexpression of a germline-specific *pgl-1::RFP* reporter in somatic cells following pulses of Myb expression under control of the *hsp-16* promoter (Figure S4). However, this promoter may not function sufficiently early during development to turn on germline genes in somatic cells ^52^. It is also possible that sustained rather than transient expression of Myb in a somatic tissue would be required to mimic this phenotype of endogenous synMuvB loss-of-function mutants ^36,37^.

The conservation in *C. elegans* of the amino acids in LIN9 and LIN52 that contact Myb in the crystal structure of their human homologs suggests that there may be another as yet unknown protein in nematodes (and perhaps other species) that can bind to the same LIN9-LIN52 structure in order to activate genes that are repressed by MuvB and DREAM complexes (Figure 7). Furthermore, the ability of *C. elegans* LIN9 and LIN52 to bind *Drosophila* Myb with a similar affinity and to discriminate among Myb mutants in a fashion similar to *Drosophila* LIN9 and LIN52 suggest a strong selective pressure during the evolution of the Myb-less Nematoda to retain the amino acids that directly contact Myb in humans and *Drosophila*.

It is possible that the amino acids in LIN9 and LIN52 that contact Myb are also essential for the structural integrity of these proteins, thus providing another explanation for their evolutionary conservation. In addition, the LIN9-LIN52 heterodimerization interface may be highly conserved because it is essential for incorporation of these proteins into the MuvB complex^45^. On the other hand, the presence of a conserved LIN9 Myb-binding domain in species that have neither an animal-type Myb protein nor a LIN52 protein suggests that this domain of LIN9 may also interact directly with other proteins (Figures 1 and 3).

Genes encoding components of the Myb-MuvB and DREAM complexes are frequently altered in human cancer. For example, 47% of a series of 2051 primary breast cancers were found to contain mutations in one of more of these genes (Figure S1). Although the *MYBL2* gene encoding the B-Myb protein is altered in only 4% of breast cancers, increased levels of expression of this gene occur more frequently, particularly in basal-like and triple-negative (ER-, PR-, HER2-) breast cancers that generally have a poor prognosis ^27,29^. Indeed, *MYBL2* is one of a small number of genes included in the Oncotype DX gene expression test that is widely used to predict clinical outcomes and plan treatment for patients with breast cancer ^28,53^.

The remarkable conservation of the Myb-binding domains of LIN9 and LIN52 described above suggests that it might be difficult to develop resistance to therapeutic drugs that target this site in order to prevent the binding of B-Myb in cancer patients. Furthermore, the conservation of a functional interaction of Myb and LIN9-LIN52 family proteins *in vivo* argues that inexpensive, genetically tractable model organisms such as flies and worms may be useful for screening for biological activity following the identification of lead compounds that inhibit binding *in vitro*.

Our studies highlight the strengths and weaknesses of using evolutionary conservation of primary sequences to predict protein function *in vitro* and *in vivo*. Database searching followed by local alignments of protein sequences permitted the identification of homologs containing conserved Myb-binding and MuvB-binding domains in highly divergent eukaryotic species (Figures 1-3). These data in turn led to inferences about selective pressures for retention of these domains during long periods of evolution. These data also led to direct tests of whether a long lost “key” (the animal-type Myb protein of *Drosophila*) is capable of opening an ancient “lock” *in vivo* (the DREAM complex of *C. elegans*) despite over 500 million years of evolution in the absence of this “key” (Figure 7). The high degree of sequence conservation in the MuvB-binding domains of vertebrate c-Myb proteins also led to the prediction that this domain would bind to the Myb-binding domain of LIN9-LIN52. However, this prediction was not borne out ^45^. This surprising result highlights the need to test hypotheses based on molecular evolution analyses by direct experimentation.

The MuvB-binding domain of vertebrate c-Myb proteins may have been conserved for another function common to all animal-type Myb proteins. Previous studies have provided evidence for negative auto-regulation of animal-type Myb proteins ^54–60^. Phosphorylation of B-Myb in and around the proline-rich hinge region by the cyclin A-CDK2 protein kinase (Figure 2) has been shown to cause a conformational change mediated by a peptidyl-prolyl isomerase that can also interact with the c-Myb protein ^61,62^. These results suggest that in addition to interacting with LIN9 and LIN52, the conserved C-terminal MuvB-binding domain may also interact directly or indirectly with the N-terminal DNA-binding domain within animal-type Myb proteins. This could provide a means to regulate the activity of these proteins via phosphorylation and isomerization of their conserved central proline-rich hinge region (Figures 1 and 2). Such a mechanism could explain the selective pressure for retaining a conserved MuvB-binding domain in c-Myb proteins that cannot bind to LIN9 and LIN52.

## MATERIALS AND METHODS

### Database Searching and Sequence Alignment

Homologous protein sequences were identified by BLASTp or tBLASTn searches of the non-redundant protein or nucleotide sequence databases at NCBI as of August 2019 [https://blast.ncbi.nlm.nih.gov/Blast.cgi] for clades of interest based on the phylogenetic trees at the Tree of Life Web Project [http://tolweb.org/tree/]. Full-length human or *Drosophila* proteins were used as query sequences. Search results with the lowest E values (using a cut-off of <1e-10 and the BLOSUM62 scoring matrix) were then “back” BLASTed against the non-redundant human or Drosophila protein databases to identify putative orthologs of the query sequences. For *Trichoplax adherens*, all genomic DNA assemblies available at ENSEMBL, some of which were not yet in the NCBI database, were also searched [https://metazoa.ensembl.org/Trichoplax_adhaerens/Info/Annotation/]. Conserved domains were identified in homologous protein families and aligned using the MACAW local alignment tool with the BLOSUM62 scoring matrix ^63^. Accession numbers for all sequences used in these alignments are provided in the Supplemental Information.

### Recombinant Protein Production and Binding Assays

LIN9 and LIN52 Myb-binding domains were co-expressed in *E. coli* ^45^. The open reading frames were synthesized as gBLOCK cassettes (IDT, Coralville, IA), cloned into the following plasmids, then verified by DNA sequencing. LIN9 (*C. elegans* residues 442-559 or *D. melanogaster* residues 571-699) was expressed from a pRSF plasmid without an affinity tag. LIN52 (*C. elegans* residues 75-139 or *D. melanogaster* residues 88-152) was expressed from an engineered pGEX plasmid with an N-terminal GST tag and a TEV protease cleavage site. Proteins were expressed overnight by addition 1 mM IPTG at 20 °C. The heterodimer complex was first purified using glutathione-sepharose affinity chromatography in a buffer containing 40 mM Tris, 150 mM NaCl, and 1 mM DTT (pH 8.0). Following elution in buffer containing 10 mM glutathione, the heterodimer was further purified with anion exchange chromatography, cleaved with TEV at 4 °C overnight, and the sample was then re-passed over glutathione-sepharose resin to remove the free GST. Final purification was achieved through Superdex 200 size-exclusion chromatography. Wild-type or mutant *D. melanogaster* Myb (residues 602-632) was expressed from the pGEX vector and purified as described for the LIN9-LIN52 complex.

For isothermal titration calorimetry experiments, proteins were concentrated as needed following purification and dialyzed overnight in the same buffer containing 20 mM Tris, 150 mM NaCl, and 1 mM beta mercaptoethanol (pH 8.0). The Myb fragment (∼500 μM) was loaded into the MicroCal VP-ITC calorimeter syringe and injected into LIN9-LIN52 heterodimer (∼25-50 μM) at 25 °C. Experiments were done in triplicate and data analyzed using the Origin ITC software package.

### Transgenic Nematode Production

Plasmids containing transgenes for expression in *C. elegans* were constructed using the Gateway system to join four different elements ^64^. The 5’ entry plasmid containing the *C. elegans hsp-16* promoter was pCM1.56. The middle entry plasmid containing a *GFP* open reading frame with *C. elegans* introns was pCM1.53. The opening reading frames encoding wild type, C-term, K606A/W607A, or D617A/Q618A mutant *Drosophila melanogster* Myb were each cloned in-frame into pCM1.53 ^11^ The 3’ entry plasmid containing the 3’ UTR of *C. elegans tbb-2* was pCM1.36. The destination vector was CFJ150, which contains an *unc-119* rescue fragment and the genomic DNA sequences flanking a *Mos1* transposon insertion on Chromosome II. The final plasmids were validated by DNA sequencing. Site-specific integration of transgenes in the host strain SS1057 with the genotype *unc-119(ed9) III* was accomplished with the CRISPR-Cas9 system ^50,51^. The single guide RNA with sequence GATATCAGTCTGTTTCGTAA targeting chromosome II near the ttTi5605 *Mos1* insertion site was cloned into pJW1219 using Q5 Site-Directed Mutagenesis (NEB, Ipswich, MA). Worms containing unintegrated transgenic DNA arrays were eliminated by screening for the absence of plasmids pCFJ90 (*Pmyo-2::mCherry)* and pCFJ104 (*Pmyo-3::mCherry)* expressing mCherry in the pharynx and body wall, respectively. Insertion sites were verified by PCR of genomic DNA (Figure S3).

Strains established from individual hermaphrodites and were maintained as homozygotes. The expression of GFP or GFP::Myb fusion proteins by integrated transgenes was verified by fluorescence microscopy four hours after a 15 minute heat shock at 37° C. Each transgene was then crossed into a *lin15-A(n767)* background. Homozygous strains were again established from individual hermaphrodites and verified by PCR of genomic DNA (Figure S3). Each transgene was also crossed into a *pgl1-1::RFP* reporter background and homozygous strains were established (Figure S4) ^65^.

### Phenotypic Analysis of Nematodes

Young adult hermaphrodites were incubated at 22^0^ C on 10cm NGM plates with an OP50 bacterial lawn. After depositing embryos for 24 hours, adults were removed with an aspirator. Approximately 24 hours later a mixed population of L2 and L3 larvae were subjected to a 15 minute heat shock in a 37° C water bath. Plates were then incubated for approximately 48 hours until all worms were adults. Worms were collected in S buffer (100 mM sodium chloride, 50 mM potassium phosphate, pH 6.0), pelleted by centrifugation, fixed in a solution of 4% paraformaldehyde in phosphate-buffered saline (pH 7.4) for 10 minutes at 22°C, washed three times in S buffer, then kept on ice prior to scoring. The presence or absence of multiple vulvae was scored by light microscopy after worms were moved to a thin agar slab on a glass slide and then flattened with a glass coverslip.

Tests for inappropriate expression of a germline gene in somatic cells were conducted in a similar fashion in worms containing both an integrated *pgl-1::RFP* reporter gene and the *hsp-16::GFP* or *hsp-16::GFP::Myb* transgene of interest. Following heat shock at 37°C for 20 minutes, recovery for 20 minutes at 22°C, and a second heat shock at 37°C for 20 minutes, worms were cultured for an additional 3 hours at 22°C and then examined for RFP and GFP fluorescence by live imaging with a Yokogawa CSUX-1 spinning disk scanner, a Nikon TE2000-E inverted microscope, a Hamamatsu ImageEM X2 camera, solid state excitation lasers at 488nm and 561nm, and 500–550nm and 573–613nm emission filters (Figure S4).

## ACKNOWLEDGEMENTS

This research was supported by NIH research grants R01CA128836 (to J.S.L.), R01CA132685 and R01GM124148 (to S.M.R.), R01GM34059 (to S.S.), and by fellowship F31CA206244 (to K.Z.G.). We thank Andy Fire, Stuart Kim, Paul Sternberg, and Anne Villeneuve and members of their laboratories and of our own laboratories for discussions; Karen Artiles for helpful advice and instruction; Dustin Updike for providing the *pgl-1::RFP* reporter; and Lajja Mani for technical support.

## SUPPLEMENTAL INFORMATION

### Accession Numbers for Sequences Used in Alignments

| Genus species | 3R Myb | LIN9 | LIN52 |
| --- | --- | --- | --- |
| *Homo sapiens* | NP_001123645.1<br>NP_001073885.1<br>NP_002457.1 | NP_775106.2 | Q52LA3.1 |
| *Branchiostoma belcheri* | XM_019789564.1 | XP_019641321.1 | XP_019625708.1 |
| *Ciona intestinalis* | XP_002119476.1 | XP_004225686.3 | XP_002124889.1 |
| *Saccoglossus kowalevskii* | XP_006814133.1 | XP_006817991.1 | XP_002734202.1 |
| *Strongylocentrotus purpuratus* | AAC47807.1 | XP_011682985.1 | XP_011662533.1 |
| *Drosophila melanogaster* | NP_511170.1 | NP_569958.2 | NP_001284908.1 |
| *Caenorhabditis elegans* | not found | NP_001023015.1 | NP_001255033.1 |
| *Priapulus caudatus* | XM_014810889 | XP_014666074.1 | XP_014662975.1 |
| *Mizuhopecten yessoensis* | XP_021368124.1 | XP_021347997.1 | XP_021378740.1 |
| *Lingula anatina* | XM_013551519 | XP_013416607.1 | XP_013388901.1 |
| *Stylophora pistillata* | XP_022808072.1 | XP_022790877.1 | XP_022785302.1 |
| *Trichoplax adherens* | RDD37379.1 | ASM15027v1:scaffold_1:<br>7596131:7601845:1 | RDD43144.1 |
| *Fusarium sp.AF-4* | RSL97542.1 | not found | not found |
| *Dictyostelium discoideum* | XP_628877.1 | XP_637607.1 | XP_643920.1 |
| *Theobroma cacao* | EOY07526.1 | XP_007035525.2 | not found |
| *Stentor coeruleus* | OMJ80050.1 | OMJ86060.1 | not found |

### Sequences of DNA Cassettes Used to Encode Recombinant Proteins

The following sequences were synthesized as double-stranded gBLOCK cassettes (IDT, Coralville, IA) and were then digested with the indicated restriction enzymes (at the underlined recognition sites), gel purified, and cloned into the bacterial expression plasmids described in the main manuscript in order to produce recombinant proteins for isothermal titration calorimetry.

Dm Mip130/LIN9 Myb-binding domain (digested with NdeI and XhoI)

~~~
CCCGGGATC<u>CATATG</u>ACGAGAAATCGCGGCTACTCCACCTCGCTGTTGGA
GCACCTGGTGCGCCTGGAGAAGTACATTGCAGTTAAGGCGGATCGAATCC
AGCGGCTCAACAAGATGAACGGCACCGCCGAGCTGGCGATGGGCGATATG
ATAAGCCATGACGAGAATGGGGATCGCCATCGCCGACAGATTGCAGTCAA
CTTCCAGCGCCAGTATGCCTTCAACATCGTGACCATCGAGCGCATCAACG
CCGAGCTCATGTTCGAGCTCACCAAGGTGCAGGAGCTGTCCTCCAGCCTG
ACACGCAATCCCAATGTCCAGGCCATGATTTCGCCGACATATTTGCGCGA
AGAGTGCCGCGCCAAGGCGTCACAGACGGTCGACGACATCAACAAGGGCA
TGTAG<u>CTCGAG</u>AGATCTCCCGGG
~~~

Ce LIN9 Myb-binding domain (digested with NdeI and XhoI)

~~~
CCCGGGATC<u>CATATG</u>GAAATGGTTGGAAATTTCCCGCTGAAATTCCTTGT
GAATCTTGTGAAACTGACGAAATTAATTGATATCAAAAAGGGATTGATAC
GACAATTGAACGAATTGAATGCGGATGCCGAGATACAAAATATGACGTCA
GACAAATATTCGAAAGCTTTTCAGGAGAAATACGCCAAAACTATCATCGA
TCTGGAACATGTGAATCAGAATATAGATATCAATATGAATGGAATTCAAG
ATCACCACATGTATTTCTCTTCGAATGATATTTCAACGTCAAATATGAAA
CCTGAAGCAGTTAGACAAATGTGCTCTCAACAAGCTGGAAGATTTGTAGA
GCACTGTAATCAAGGATTATAGCTCGAGAGATCTCCCGGG
~~~

Dm LIN52 Myb-binding domain (digested with EcoRI and XhoI)

~~~
CCCGGGATCCGG<u>GAATTC</u>ACGCGCAGCACCAACTACACCAGCAATCTGAC
CGATGATGATATGGTTAAGATTAATGAACTAGCCCAGCTCCCTCCCGAGG
ATCTGATCGATAAAATAAAGTCAATGCATGATGAAATTTACCAGCTGGGA
CTGCGTGAGGCAATGGAGATGACTCGTGGGAAACTGCTGGGCATCTTTGA
CCGGGATCGCGCTTAG<u>CTCGAG</u>AGATCTCCCGGG
~~~

Ce LIN52 Myb-binding domain (digested with EcoRI and XhoI)

~~~
CCCGGGATCCGG<u>GAATTC</u>TACGAATCGCCATACAAGAATATTTCGTTTCT
CAAGGAAGATGCTGTGACTGTTAATACAATGAGCCACTGCCCAGCCGACG
ATATCGCCAAGCTCATCCGAAACATTCAAAACTCGGTGTACACTCTTGGA
ATCGAAGAAGCTCGCCAGTGCCGACGTGGAAAGTTGCTCAACGTGCTGAA
ACCCACTGGCTCGTAG<u>CTCGAG</u>AGATCTCCCGGG
~~~

WT Dm Myb MuvB-binding domain (digested with EcoRI and XhoI)

~~~
CCCGGGATCCGG<u>GAATTC</u>GTCATTGATCCCAAGTGGGCACGCGTCGCTTG
TGGCAAGTCCAGAGATCAAATGTTTATGGAGGAGCAGGCTTATGCGTGCC
TCAAAAATCTGTAG<u>CTCGAG</u>AGATCTCCCGGG
~~~

KW Dm Myb MuvB-binding domain (digested with EcoRI and XhoI)

~~~
CCCGGGATCCGG<u>GAATTC</u>GTCATTGATCCCGCGGCGGCACGCGTCGCTTG
TGGCAAGTCCAGAGATCAAATGTTTATGGAGGAGCAGGCTTATGCGTGCC
TCAAAAATCTGTAG<u>CTCGAG</u>AGATCTCCCGGG
~~~

DQ Dm Myb MuvB-binding domain (digested with EcoRI and XhoI)

~~~
CCCGGGATCCGG<u>GAATTC</u>GTCATTGATCCCAAGTGGGCACGCGTCGCTTG
TGGCAAGTCTAGAGCTGCAATGTTTATGGAGGAGCAGGCTTATGCGTGCC
TCAAAAATCTGTAG<u>CTCGAG</u>AGATCTCCCGGG
~~~

### Sequences of GFP::Myb Junctions in C. elegans Transgenes

**WT, KW, and DQ**

~~~
GCT GCT GGG ATT ACA CAT GGC ATG GAC GAA CTA TAC AAA AGC CCA CAA
<u>ala ala gly ile thr his gly met asp glu leu tyr lys</u> ser pro gln
                   GFP
~~~

~~~
AAG CTT AAG ATG GCA AGT GCG AGC ACT GAA AAC GGC GAG GAG CTG ATG
lys leu lys <u>met ala ser ala ser thr glu gln gly glu glu leu met</u>
                                 MYB
~~~

**C-term**

~~~
GCT GCT GGG ATT ACA CAT GGC ATG GAC GAA CTA TAC AAA AGC CCA CAA
<u>ala ala gly ile thr his gly met asp glu leu tyr lys</u> ser pro gln
                   GFP
~~~

~~~
AAG CTT AAG ATG TCT GGT AGC GAT TTG AAG AGT TCG CGA ACC CAT CTC
lys leu lys Met <u>ser gly ser asp leu lys ser ser arg thr his leu</u>
                                      MYB
~~~

### Supplemental Figures

**Figure S1.**
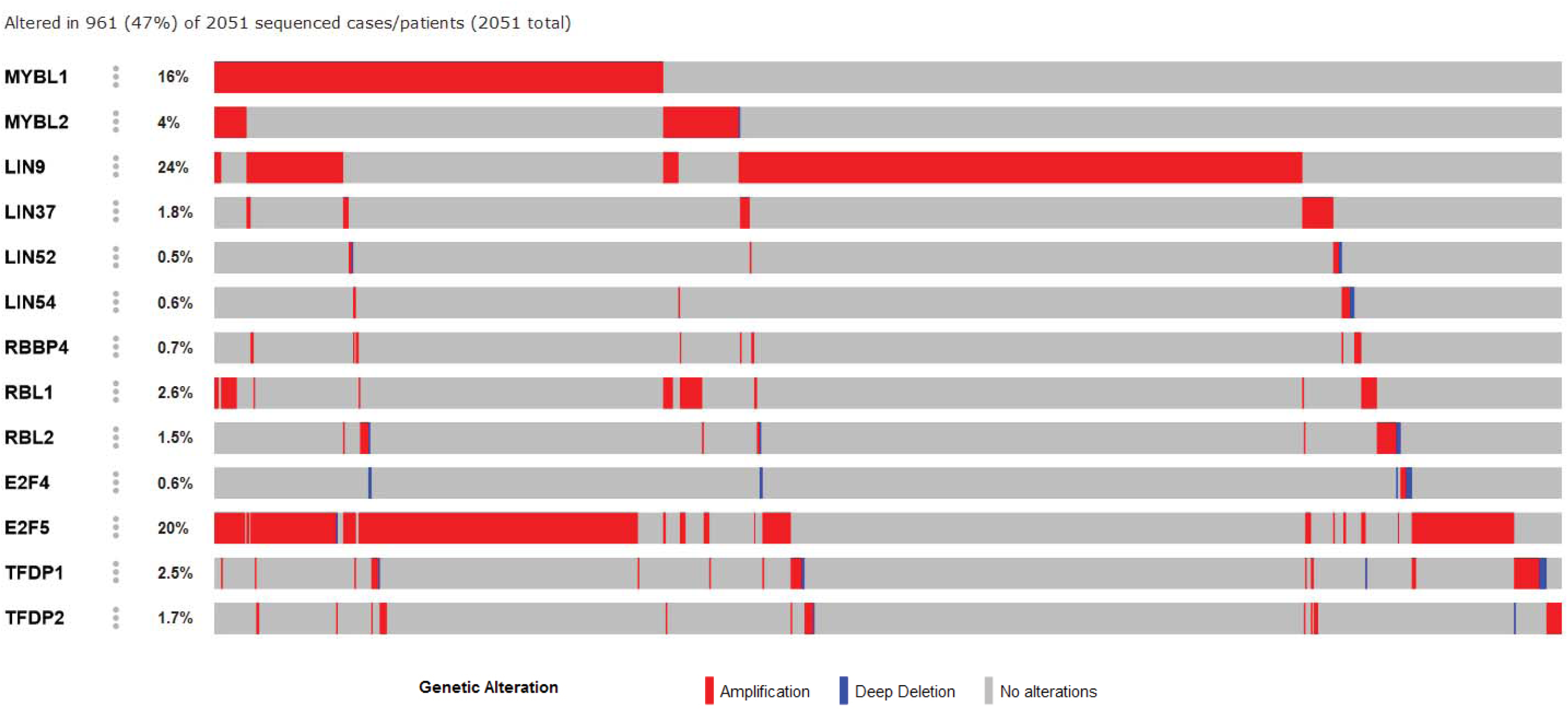
Genes Encoding the Myb-MuvB and DREAM Complex Proteins Are Frequently Altered in Human Breast Cancer. Publicly available cancer genome data from the METABRIC study were analyzed using the cBioPortal website [http://www.cbioportal.org/] ^1,2^.

**Figure S2.**
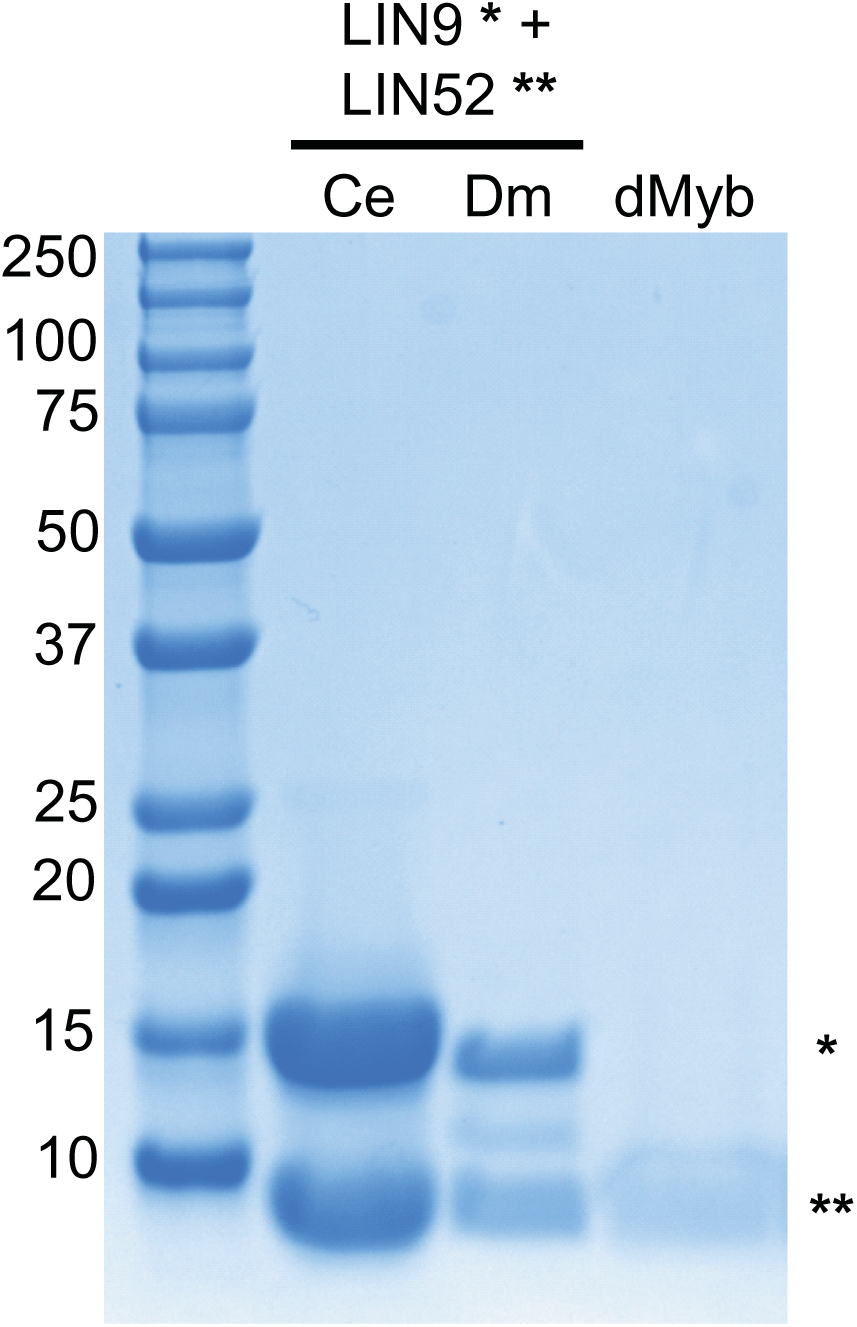
Proteins Used for Isothermal Calorimetry Studies. Coomassie-stained SDS-PAGE gel shows purified wild-type LIN9-LIN52 proteins (Ce: *C. elegans*, Dm: *D. melanogaster*) and purified dMyb.

**Figure S3.**
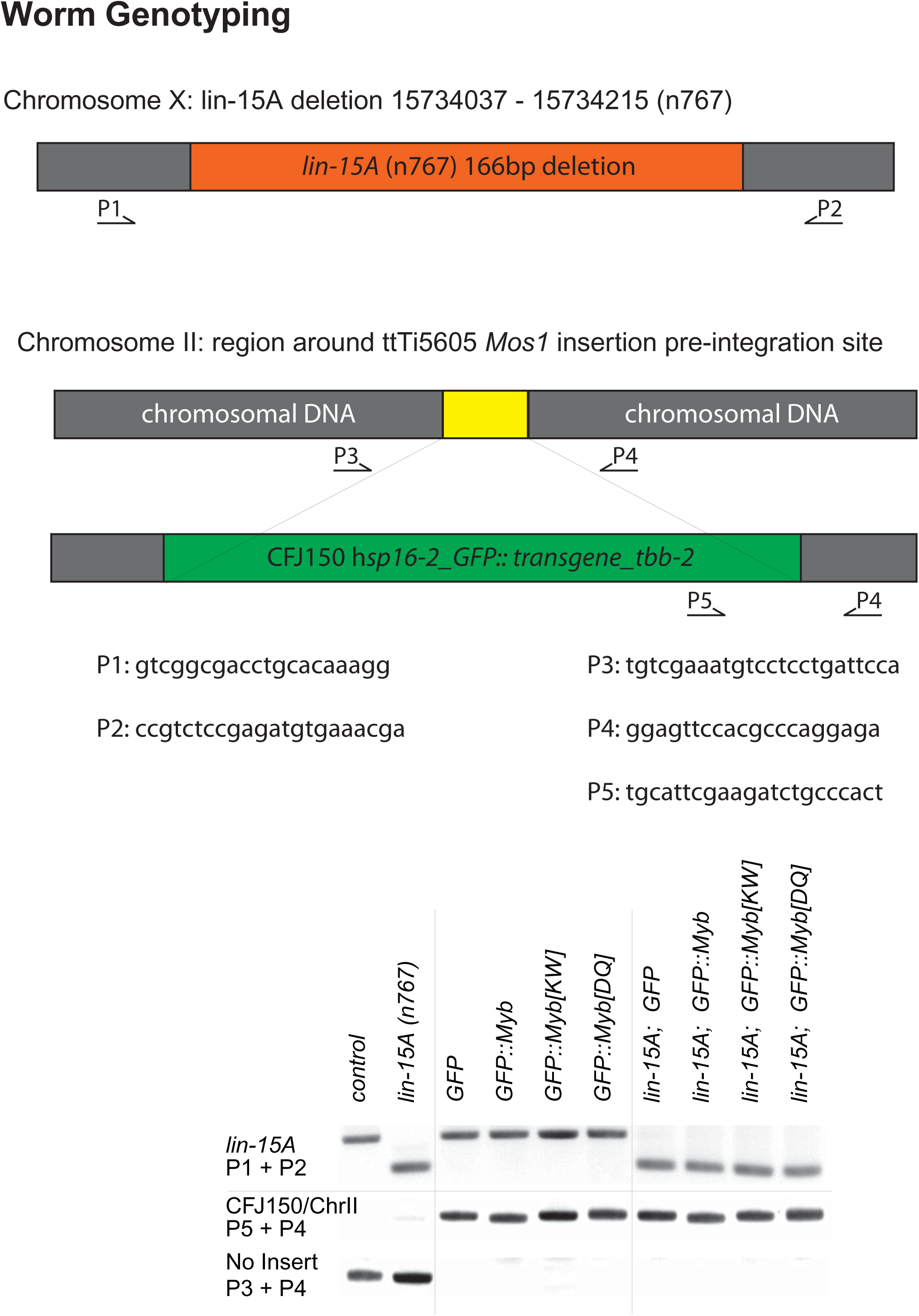
PCR Genotyping of *C. elegans* Strains. Diagrams of the loci of interest, PCR primers, and predicted amplicons are shown in the top panel. The bottom panel shows a representative PCR genotyping of strains used in this paper.

**Figure S4.**
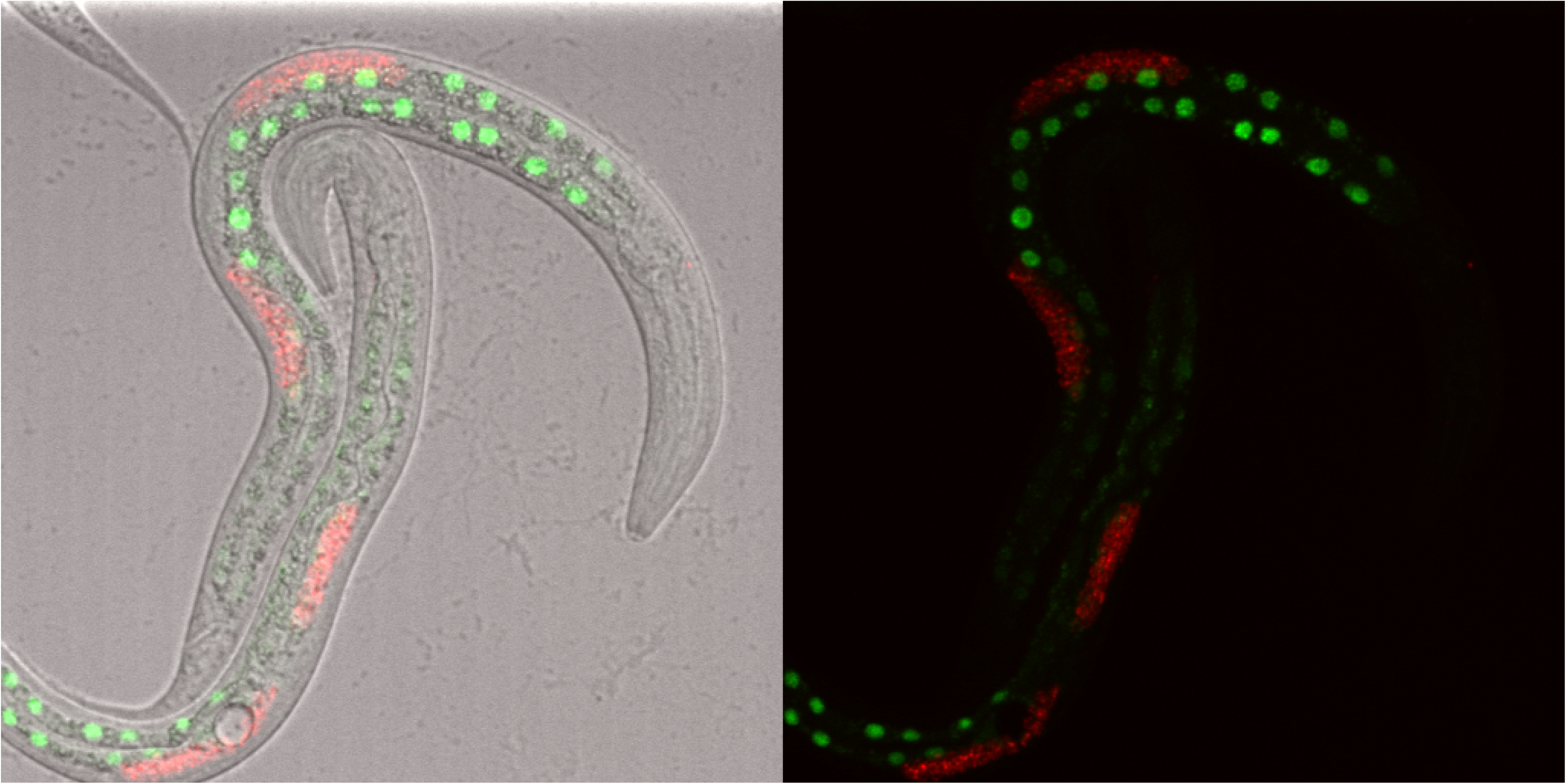
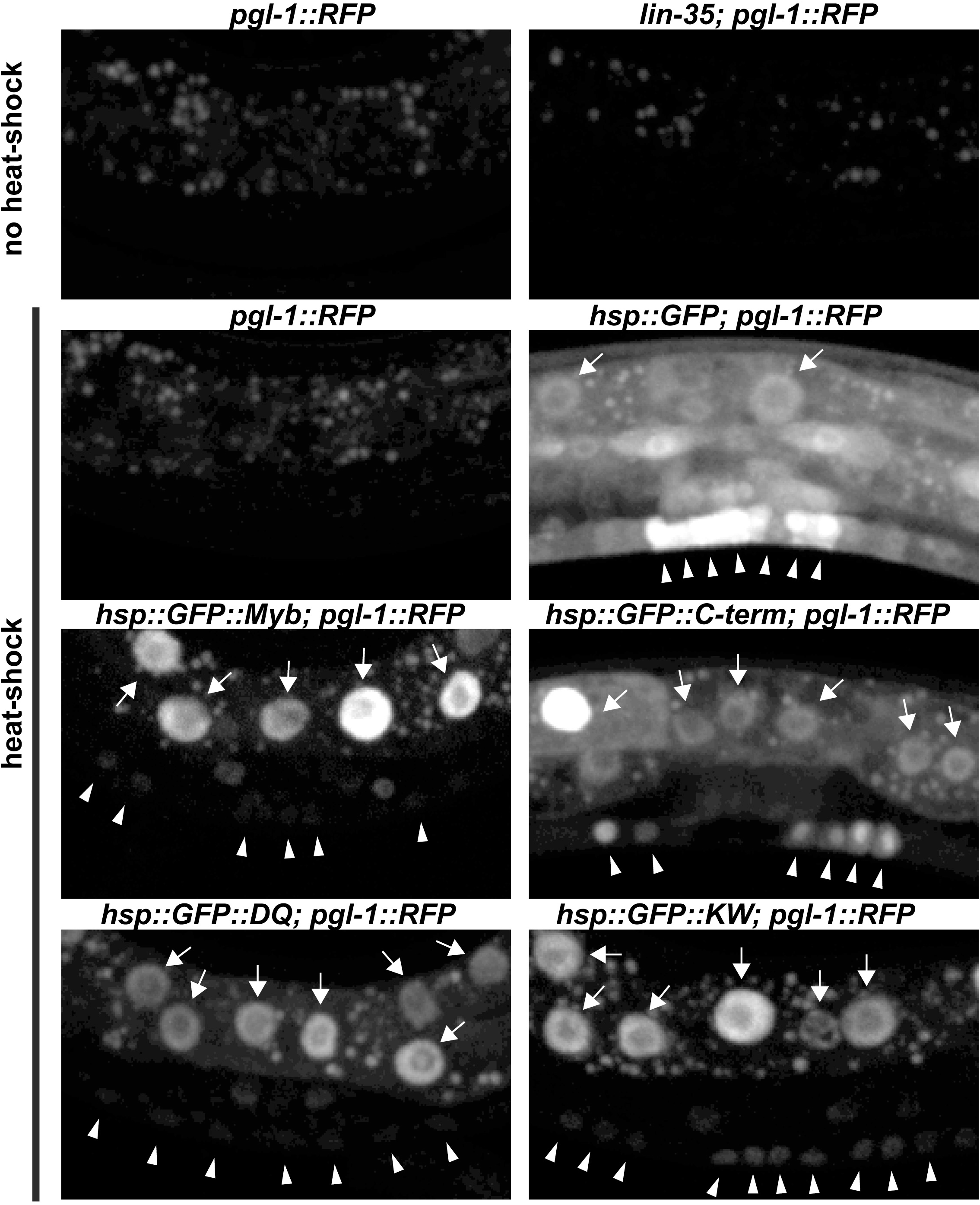
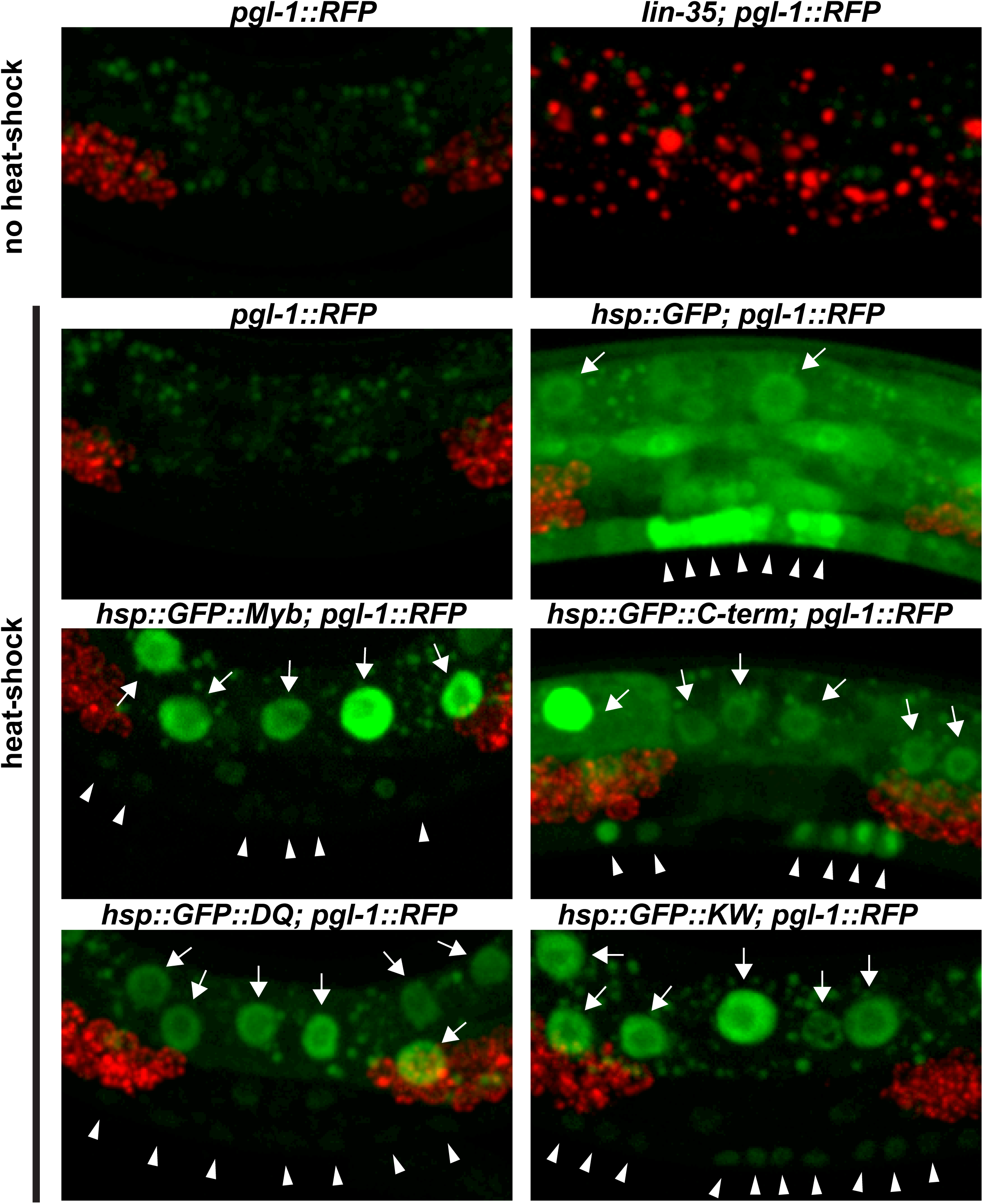
Expression of GFP and GFP::Myb Fusion Proteins in Transgenic Worms. Panel A: Photomicrographs of heat-shocked mid-L3 larvae containing an *hsp-16::GFP::Myb* transgene and a *pgl-1::RFP* reporter gene taken with a 20X objective. The left side shows GFP and RFP fluorescence superimposed on a bright-field DIC image. The right side shows only the GFP and RFP fluorescence. Panel B: Higher maginifcation images of heat-shocked mid-L3 larvae of the indicated genotypes showing only the GFP fluorescence. The punctate fluorescence observed in animals without heat shock is due to autofluorescent granules within gut cells. GFP-positive gut cell nuclei are indicated by arrows. GFP-positive nuclei of VPCs and adjacent hypodermal cells are inidicated by arrowheads. Panel C: Same microscopic fields shown in Panel B with both GFP (green) and RFP (red) fluorescence. Note the inappropriate expression of the germline *pgl-1::RFP* reporter in somatic gut cells in the *lin-35(n745)* mutant positive control, but not in the transgenic animals expressing GFP or GFP::Myb fusion proteins.

## Notes

#### Summary of Updates

Supplemental Figure S4 is new, as is the text describing it.

## LITERATURE CITED

1 Lipsick, J. S. & Wang, D. M. Transformation by v-Myb. Oncogene 18, 3047–3055 (1999).

2 Lipsick, J. S. One billion years of Myb. Oncogene 13, 223–235 (1996).

3 Davidson, C., Ray, E. & Lipsick, J. in Myb Transcription Factors: Their Role in Growth, Differentiation and Disease (ed J. Frampton) 1–33 (Kluwer Academic Publishers, 2004).

4 Katzen, A. L., Kornberg, T. B. & Bishop, J. M. Isolation of the proto-oncogene c-myb from D. melanogaster. Cell 41, 449–456 (1985).

5 Katzen, A. L. & Bishop, J. M. myb provides an essential function during Drosophila development. Proc Natl Acad Sci U S A 93, 13955–13960 (1996).

6 Manak, J. R., Mitiku, N. & Lipsick, J. S. Mutation of the Drosophila homologue of the Myb protooncogene causes genomic instability. Proc Natl Acad Sci U S A 99, 7438–7443 (2002).

7 Okada, M., Akimaru, H., Hou, D. X., Takahashi, T. & Ishii, S. Myb controls G(2)/M progression by inducing cyclin B expression in the Drosophila eye imaginal disc. Embo J 21, 675–684. (2002).

8 Davidson, C. J., Guthrie, E. E. & Lipsick, J. S. Duplication and maintenance of the Myb genes of vertebrate animals. Biol Open 2, 101–110, doi:10.1242/bio.20123152 (2013).

9 Davidson, C., Tirouvanziam, R., Herzenberg, L. & Lipsick, J. Functional Evolution of the Vertebrate Myb Gene Family: B-Myb, but neither A-Myb nor c-Myb, complements Drosophila Myb in Hemocytes. Genetics 169, 215–229 (2005).

10 Wen, H., Andrejka, L., Ashton, J., Karess, R. & Lipsick, J. S. Epigenetic regulation of gene expression by Drosophila Myb and E2F2-RBF via the Myb-MuvB/dREAM complex. Genes Dev 22, 601–614, doi:10.1101/gad.1626308 (2008).

11 Andrejka, L., Wen, H., Ashton, J., Grant, M., Iori, K., Wang, A., Manak, J. R. & Lipsick, J. S. Animal-specific C-terminal domain links myeloblastosis oncoprotein (Myb) to an ancient repressor complex. Proc Natl Acad Sci U S A 108, 17438–17443, doi:10.1073/pnas.1111855108 (2011).

12 Beall, E. L., Manak, J. R., Zhou, S., Bell, M., Lipsick, J. S. & Botchan, M. R. Role for a Drosophila Myb-containing protein complex in site-specific DNA replication. Nature 420, 833–837 (2002).

13 Sadasivam, S. & DeCaprio, J. A. The DREAM complex: master coordinator of cell cycle-dependent gene expression. Nat Rev Cancer 13, 585–595, doi:10.1038/nrc3556 (2013).

14 Lewis, P. W., Beall, E. L., Fleischer, T. C., Georlette, D., Link, A. J. & Botchan, M. R. Identification of a Drosophila Myb-E2F2/RBF transcriptional repressor complex. Genes Dev 18, 2929–2940 (2004).

15 Korenjak, M., Taylor-Harding, B., Binne, U. K., Satterlee, J. S., Stevaux, O., Aasland, R., White-Cooper, H., Dyson, N. & Brehm, A. Native E2F/RBF Complexes Contain Myb-Interacting Proteins and Repress Transcription of Developmentally Controlled E2F Target Genes. Cell 119, 181–193 (2004).

16 Katzen, A. L., Jackson, J., Harmon, B. P., Fung, S. M., Ramsay, G. & Bishop, J. M. Drosophila myb is required for the G2/M transition and maintenance of diploidy. Genes Dev 12, 831–843 (1998).

17 Fung, S. M., Ramsay, G. & Katzen, A. L. Mutations in Drosophila myb lead to centrosome amplification and genomic instability. Development 129, 347–359. (2002).

18 Manak, J. R., Wen, H., Tran, V., Andrejka, L. & Lipsick, J. S. Loss of Drosophila Myb interrupts the progression of chromosome condensation. Nat Cell Biol 9, 581–587 (2007).

19 DeBruhl, H., Wen, H. & Lipsick, J. S. The complex containing Drosophila Myb and RB/E2F2 regulates cytokinesis in a histone H2Av-dependent manner. Mol Cell Biol 33, 1809–1818, doi:10.1128/MCB.01401-12 (2013).

20 Dimova, D. K., Stevaux, O., Frolov, M. V. & Dyson, N. J. Cell cycle-dependent and cell cycle-independent control of transcription by the Drosophila E2F/RB pathway. Genes Dev 17, 2308–2320 (2003).

21 Georlette, D. et al. Genomic profiling and expression studies reveal both positive and negative activities for the Drosophila Myb MuvB/dREAM complex in proliferating cells. Genes Dev 21, 2880–2896 (2007).

22 Litovchick, L. et al. Evolutionarily Conserved Multisubunit RBL2/p130 and E2F4 Protein Complex Represses Human Cell Cycle-Dependent Genes in Quiescence. Mol Cell 26, 539–551 (2007).

23 Osterloh, L., von Eyss, B., Schmit, F., Rein, L., Hubner, D., Samans, B., Hauser, S. & Gaubatz, S. The human synMuv-like protein LIN-9 is required for transcription of G2/M genes and for entry into mitosis. Embo J 26, 144–157 (2007).

24 Pilkinton, M., Sandoval, R., Song, J., Ness, S. A. & Colamonici, O. R. Mip/LIN-9 regulates the expression of B-Myb and the induction of cyclin A, cyclin B, and CDK1. J Biol Chem 282, 168–175 (2007).

25 Muller, G. A., Stangner, K., Schmitt, T., Wintsche, A. & Engeland, K. Timing of transcription during the cell cycle: Protein complexes binding to E2F, E2F/CLE, CDE/CHR, or CHR promoter elements define early and late cell cycle gene expression. Oncotarget 8, 97736–97748, doi:10.18632/oncotarget.10888 (2017).

26 Fischer, M. & Muller, G. A. Cell cycle transcription control: DREAM/MuvB and RB-E2F complexes. Crit Rev Biochem Mol Biol 52, 638–662, doi:10.1080/10409238.2017.1360836 (2017).

27 Amatschek, S. et al. Tissue-wide expression profiling using cDNA subtraction and microarrays to identify tumor-specific genes. Cancer Res 64, 844–856 (2004).

28 Paik, S. et al. A multigene assay to predict recurrence of tamoxifen-treated, node-negative breast cancer. N Engl J Med 351, 2817–2826 (2004).

29 Thorner, A. R., Hoadley, K. A., Parker, J. S., Winkel, S., Millikan, R. C. & Perou, C. M. In vitro and in vivo analysis of B-Myb in basal-like breast cancer. Oncogene 28, 742–751 (2009).

30 Lipsick, J. S. synMuv verite--Myb comes into focus. Genes Dev 18, 2837–2844 (2004).

31 Sternberg, P. W. & Han, M. Genetics of RAS signaling in C. elegans. Trends Genet 14, 466–472 (1998).

32 Horvitz, H. R. & Sulston, J. E. Isolation and genetic characterization of cell-lineage mutants of the nematode Caenorhabditis elegans. Genetics 96, 435–454 (1980).

33 Ferguson, E. L. & Horvitz, H. R. The multivulva phenotype of certain Caenorhabditis elegans mutants results from defects in two functionally redundant pathways. Genetics 123, 109–121 (1989).

34 Cui, M., Chen, J., Myers, T. R., Hwang, B. J., Sternberg, P. W., Greenwald, I. & Han, M. SynMuv genes redundantly inhibit lin-3/EGF expression to prevent inappropriate vulval induction in C. elegans. Dev Cell 10, 667–672, doi:10.1016/j.devcel.2006.04.001 (2006).

35 Saffer, A. M., Kim, D. H., van Oudenaarden, A. & Horvitz, H. R. The Caenorhabditis elegans synthetic multivulva genes prevent ras pathway activation by tightly repressing global ectopic expression of lin-3 EGF. PLoS Genet 7, e1002418, doi:10.1371/journal.pgen.1002418 (2011).

36 Wang, D., Kennedy, S., Conte, D., Jr., Kim, J. K., Gabel, H. W., Kamath, R. S., Mello, C. C. & Ruvkun, G. Somatic misexpression of germline P granules and enhanced RNA interference in retinoblastoma pathway mutants. Nature 436, 593–597 (2005).

37 Petrella, L. N., Wang, W., Spike, C. A., Rechtsteiner, A., Reinke, V. & Strome, S. synMuv B proteins antagonize germline fate in the intestine and ensure C. elegans survival. Development 138, 1069–1079, doi:10.1242/dev.059501 (2011).

38 Tabuchi, T. M., Rechtsteiner, A., Strome, S. & Hagstrom, K. A. Opposing activities of DRM and MES-4 tune gene expression and X-chromosome repression in Caenorhabditis elegans germ cells. G3 (Bethesda) 4, 143–153, doi:10.1534/g3.113.007849 (2014).

39 Boxem, M. & van den Heuvel, S. C. elegans class B synthetic multivulva genes act in G(1) regulation. Curr Biol 12, 906–911 (2002).

40 Hsieh, J., Liu, J., Kostas, S. A., Chang, C., Sternberg, P. W. & Fire, A. The RING finger/B-box factor TAM-1 and a retinoblastoma-like protein LIN-35 modulate context-dependent gene silencing in Caenorhabditis elegans. Genes Dev 13, 2958–2970 (1999).

41 Sim, C. K., Perry, S., Tharadra, S. K., Lipsick, J. S. & Ray, A. Epigenetic regulation of olfactory receptor gene expression by the Myb-MuvB/dREAM complex. Genes Dev 26, 2483–2498, doi:10.1101/gad.201665.112 (2012).

42 Beall, E. L., Bell, M., Georlette, D. & Botchan, M. R. Dm-myb mutant lethality in Drosophila is dependent upon mip130: positive and negative regulation of DNA replication. Genes Dev 18, 1667–1680 (2004).

43 Rovani, M. K., Brachmann, C. B., Ramsay, G. & Katzen, A. L. The dREAM/Myb-MuvB complex and Grim are key regulators of the programmed death of neural precursor cells at the Drosophila posterior wing margin. Dev Biol 372, 88–102, doi:10.1016/j.ydbio.2012.08.022 (2012).

44 Goetsch, P. D., Garrigues, J. M. & Strome, S. Loss of the Caenorhabditis elegans pocket protein LIN-35 reveals MuvB’s innate function as the repressor of DREAM target genes. PLoS Genet 13, e1007088, doi:10.1371/journal.pgen.1007088 (2017).

45 Guiley, K. Z., Iness, A. N., Saini, S., Tripathi, S., Lipsick, J. S., Litovchick, L. & Rubin, S. M. Structural mechanism of Myb-MuvB assembly. Proc Natl Acad Sci U S A 115, 10016–10021, doi:10.1073/pnas.1808136115 (2018).

46 Coghlan, A. Nematode genome evolution. WormBook, 1–15, doi:10.1895/wormbook.1.15.1 (2005).

47 Telford, M. J. & Copley, R. R. Animal phylogeny: fatal attraction. Curr Biol 15, R296–299, doi:10.1016/j.cub.2005.04.001 (2005).

48 Holton, T. A. & Pisani, D. Deep genomic-scale analyses of the metazoa reject Coelomata: evidence from single- and multigene families analyzed under a supertree and supermatrix paradigm. Genome Biol Evol 2, 310–324, doi:10.1093/gbe/evq016 (2010).

49 White-Cooper, H., Leroy, D., MacQueen, A. & Fuller, M. T. Transcription of meiotic cell cycle and terminal differentiation genes depends on a conserved chromatin associated protein, whose nuclear localisation is regulated. Development 127, 5463–5473 (2000).

50 Dickinson, D. J., Ward, J. D., Reiner, D. J. & Goldstein, B. Engineering the Caenorhabditis elegans genome using Cas9-triggered homologous recombination. Nat Methods 10, 1028–1034, doi:10.1038/nmeth.2641 (2013).

51 Frokjaer-Jensen, C., Davis, M. W., Hopkins, C. E., Newman, B. J., Thummel, J. M., Olesen, S. P., Grunnet, M. & Jorgensen, E. M. Single-copy insertion of transgenes in Caenorhabditis elegans. Nat Genet 40, 1375–1383, doi:10.1038/ng.248 (2008).

52 Stringham, E. G., Dixon, D. K., Jones, D. & Candido, E. P. Temporal and spatial expression patterns of the small heat shock (hsp16) genes in transgenic Caenorhabditis elegans. Mol Biol Cell 3, 221–233, doi:10.1091/mbc.3.2.221 (1992).

53 Bhutiani, N., Vuong, B., Egger, M. E., Eldredge-Hindy, H., McMasters, K. M. & Ajkay, N. Evaluating patterns of utilization of gene signature panels and impact on treatment patterns in patients with ductal carcinoma in situ of the breast. Surgery, doi:10.1016/j.surg.2019.04.044 (2019).

54 Sakura, H., Kanei-Ishii, C., Nagase, T., Nakagoshi, H., Gonda, T. J. & Ishii, S. Delineation of three functional domains of the transcriptional activator encoded by the c-myb protooncogene. Proc Natl Acad Sci U S A 86, 5758–5762 (1989).

55 Ramsay, R. G., Ishii, S. & Gonda, T. J. Increase in specific DNA binding by carboxyl truncation suggests a mechanism for activation of Myb. Oncogene 6, 1875–1879 (1991).

56 Dubendorff, J. W., Whittaker, L. J., Eltman, J. T. & Lipsick, J. S. Carboxy-terminal elements of c-Myb negatively regulate transcriptional activation in cis and in trans. Genes Dev 6, 2524–2535 (1992).

57 Dash, A. B., Orrico, F. C. & Ness, S. A. The EVES motif mediates both intermolecular and intramolecular regulation of c-Myb. Genes Dev 10, 1858–1869 (1996).

58 Ansieau, S., Kowenz-Leutz, E., Dechend, R. & Leutz, A. B-Myb, a repressed trans-activating protein. J Mol Med 75, 815–819 (1997).

59 Lane, S., Farlie, P. & Watson, R. B-Myb function can be markedly enhanced by cyclin A-dependent kinase and protein truncation. Oncogene 14, 2445–2453 (1997).

60 Dubendorff, J. W. & Lipsick, J. S. Transcriptional regulation by the carboxyl terminus of c-Myb depends upon both the Myb DNA-binding domain and the DNA recognition site. Oncogene 18, 3452–3460 (1999).

61 Leverson, J. D. & Ness, S. A. Point mutations in v-Myb disrupt a cyclophilin-catalyzed negative regulatory mechanism. Mol Cell 1, 203–211 (1998).

62 Werwein, E., Cibis, H., Hess, D. & Klempnauer, K. H. Activation of the oncogenic transcription factor B-Myb via multisite phosphorylation and prolyl cis/trans isomerization. Nucleic Acids Res 47, 103–121, doi:10.1093/nar/gky935 (2019).

63 Schuler, G. D., Altschul, S. F. & Lipman, D. J. A workbench for multiple alignment construction and analysis. Proteins Struct.Funct.Genet. 9, 180–190 (1991).

64 Merritt, C. & Seydoux, G. Transgenic solutions for the germline. WormBook, 1–21, doi:10.1895/wormbook.1.148.1 (2010).

65 Marnik, E. A., Fuqua, J. H., Sharp, C. S., Rochester, J. D., Xu, E. L., Holbrook, S. E. & Updike, D. L. Germline Maintenance Through the Multifaceted Activities of GLH/Vasa in Caenorhabditis elegans P Granules. Genetics, doi:10.1534/genetics.119.302670 (2019).

66 Guiley, K. Z., Liban, T. J., Felthousen, J. G., Ramanan, P., Litovchick, L. & Rubin, S. M. Structural mechanisms of DREAM complex assembly and regulation. Genes Dev 29, 961–974, doi:10.1101/gad.257568.114 (2015).

67 Webb, B. & Sali, A. Protein Structure Modeling with MODELLER. Methods Mol Biol 1654, 39–54, doi:10.1007/978-1-4939-7231-9_4 (2017).

## Literature Cited

1. Gao, J. et al. Integrative analysis of complex cancer genomics and clinical profiles using the cBioPortal. Sci Signal 6, pl1, doi:10.1126/scisignal.2004088 (2013).

2. Pereira, B. et al. The somatic mutation profiles of 2,433 breast cancers refines their genomic and transcriptomic landscapes. Nat Commun 7, 11479, doi:10.1038/ncomms11479 (2016).

